# Bacterial DNA methylation amplifies the shared A-to-G and G-to-A substitution bias in nanopore sequencing

**DOI:** 10.64898/2026.01.14.699587

**Authors:** Xudong Liu, Qiutao Ding, Yanwen Shao, Zhihao Guo, Ying Ni, Lu Fan, Yi Yang, Kaixin Chen, Mengsu Yang, Runsheng Li

## Abstract

Nanopore sequencing enables direct analysis of native DNA modifications, but these modifications can also introduce systematic basecalling errors. Here, we investigated how bacterial DNA methylation contributes to nanopore substitution errors using matched native whole-genome sequencing (WGS) and methylation-depleted whole-genome amplification (WGA) datasets from six bacterial species across CycloneSEQ (CS) and Oxford Nanopore Technologies platforms (ONT). Adenine-to-guanine (A2G) and guanine-to-adenine (G2A) substitutions were consistently enriched across species, nanopore platforms, sequencing configurations, and basecalling models, indicating a shared nanopore-associated substitution bias. This enrichment remained detectable in WGA reads, whereas native WGS reads showed a further increase. Using independent PacBio methylation profiling, we found that the WGS-associated substitution excess was strongly concentrated around bacterial methylation motifs and was minimal in non-motif regions. The magnitude and substitution pattern of these errors varied among motifs and sequencing platforms. At a subset of motif-associated positions, alternative-base fractions exceeded 50%, producing SNP-like signals. In ONT data, 28 WGS-specific calls survived the applied SNP-calling filters, all within PacBio-supported methylation contexts, and most showed significant strand imbalance. Together, these results identify a shared A2G/G2A substitution bias in all nanopore datasets and support a methylation-associated increase in these errors near bacterial methylation motifs, with some loci producing SNP-like artifacts.

## Introduction

Nanopore sequencing enables direct analysis of native DNA while providing long reads that are increasingly used for bacterial genome assembly and epigenetic profiling (Tourancheau et al. 2021; Wang et al. 2021; Sereika et al. 2022). Unlike sequencing-by-synthesis technologies, nanopore sequencing infers nucleotide sequences from changes in ionic current as DNA passes through a nanopore, allowing DNA modifications to be detected without additional conversion or enrichment steps (Deamer et al. 2016; Rand et al. 2017; Simpson et al. 2017). This capability has enabled genome-wide profiling of bacterial N6-methyladenine (6mA), N4-methylcytosine (4mC), and 5-methylcytosine (5mC), as well as *de novo* identification of their associated sequence motifs (Tourancheau et al. 2021; Kulkarni et al. 2026). However, because modified nucleotides also alter the current signal used for canonical basecalling, modifications in native DNA can complicate sequence determination and introduce basecalling errors.

Nanopore sequencing errors are not uniformly distributed across sequence contexts or substitution types. Previous analyses have identified recurrent sequence-dependent errors and an excess of transition substitutions, including confusion between adenine and guanine (Delahaye and Nicolas 2021). Improvements in pore chemistry and basecalling models have substantially increased overall nanopore read accuracy (Sereika et al. 2022; Ni et al. 2023; Sanderson et al. 2024), but systematic errors remain important because the same incorrect base can repeatedly occur at the same genomic position across independent reads. Whether recurrent substitution biases represent shared features of nanopore sequencing or are mainly dependent on individual platforms or basecalling models remains incompletely understood.

Bacterial DNA methylation is one established source of systematic nanopore errors. Whole-genome amplification, which strongly depletes native DNA modifications, can reduce modification-associated nanopore errors and improve bacterial genome accuracy (Chiou et al. 2023). Our previous study showed that bacterial DNA modifications can generate strongly strand-biased nanopore substitutions near methylation motifs and that these errors can be used for de novo methylation discovery (Liu et al. 2024a). Other studies have linked methylation-associated basecalling errors to inaccurate bacterial genotyping and outbreak reconstruction and have shown that amplification-derived DNA can reduce many of these errors (Dabernig-Heinz et al. 2024; Lohde et al. 2024).

Despite these advances, two questions remain insufficiently characterized. First, it remains unclear whether particular substitution biases are shared across different nanopore sequencing platforms and persist after depletion of native DNA methylation. Second, the extent to which bacterial methylation further elevates these recurrent substitution types has not been systematically assessed across platforms. Addressing these questions requires matched comparisons of native and amplified DNA, together with independent identification of methylation contexts, to distinguish the background substitution pattern from the additional errors associated with native DNA methylation.

Here, we analyzed matched native WGS and methylation-depleted WGA datasets from six bacterial species using CS and ONT R10.4.1 (R10). We compared substitution spectra across nanopore platforms, sequencing configurations, and basecalling models, with PacBio reads providing a comparison from a different sequencing technology. We then used paired WGS-minus-WGA differences and independent PacBio methylation profiling to examine how native DNA methylation was associated with changes in these spectra. Our results identify A2G and G2A enrichment as a recurrent feature of the tested nanopore datasets that persists after amplification. Native DNA shows additional elevation of these substitutions around methylation motifs, with effects that vary among motifs and platforms. We further examined high-fraction error sites and SNP calls to illustrate the downstream consequences of this pattern.

## Results

### Paired WGS and WGA datasets enable controlled analysis of methylation-associated effects

To assess how bacterial DNA methylation influences nanopore read errors, we analyzed six phylogenetically diverse bacterial strains: *Acinetobacter pittii* (Api), *Escherichia coli* (Eco), *Enterococcus faecium* (Efa), *Klebsiella pneumoniae* (Kpn), *Pseudomonas aeruginosa* (Pae), and *Salmonella enterica* (Sen). For each strain, native WGS and WGA DNA libraries were sequenced on the CS platform, with WGA serving as a matched methylation-depleted control (**Fig. 1A**). Matched R10 datasets generated from the same strains were included for cross-platform comparison. The reference genomes spanned 2.8-6.4 Mb in size and 38.0-66.4% in GC content, enabling evaluation across diverse genomic contexts (**Table S1**).

**Figure 1.**
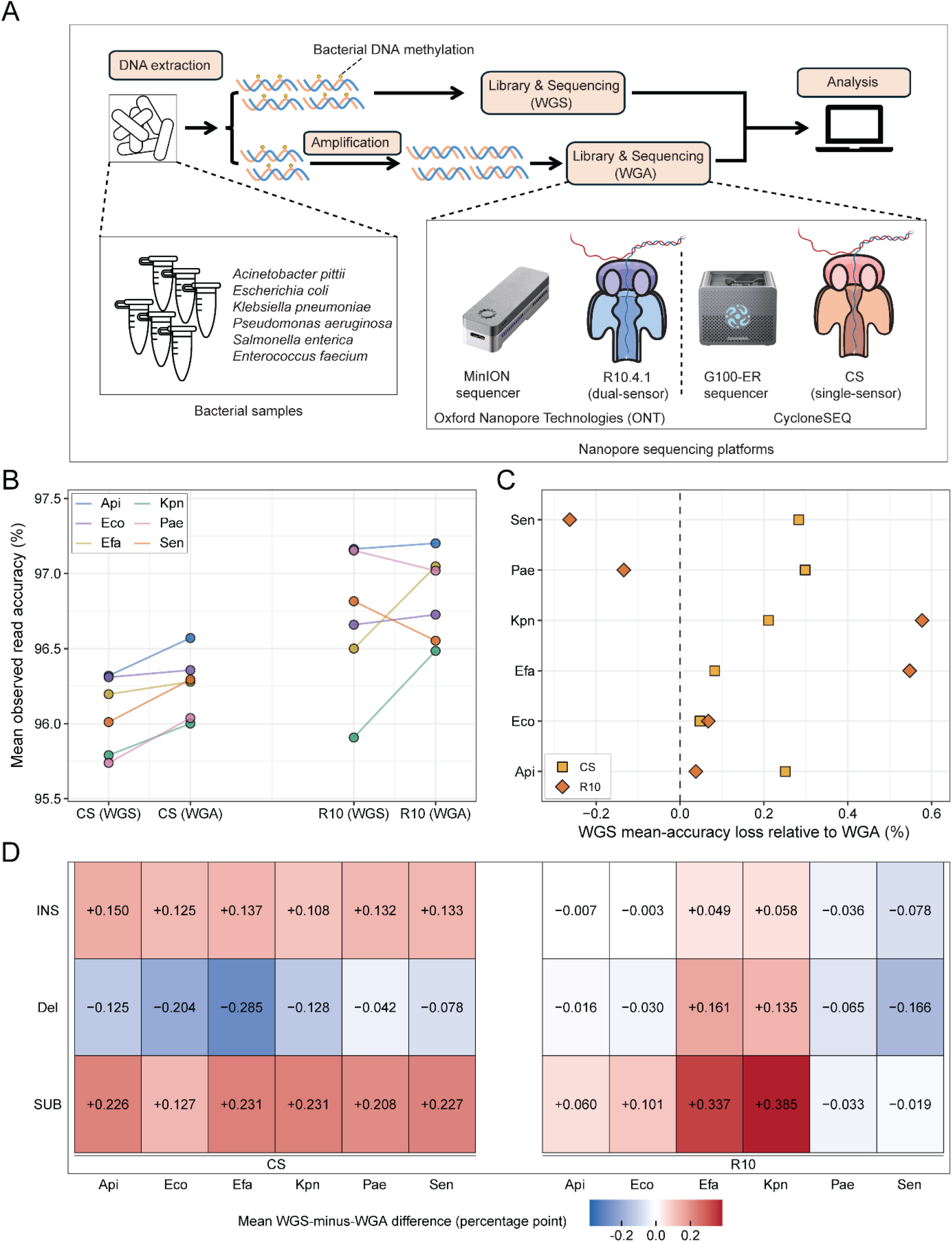
Comparison of read accuracy and error profiles between native WGS and WGA nanopore reads. **A.** Overview of the experimental design. DNA from six bacterial species was analyzed using paired native whole-genome sequencing (WGS) and whole-genome-amplified (WGA) libraries. WGA was used as a methylation-depleted control for comparison with native DNA. Libraries were sequenced using CycloneSEQ (CS) and Oxford Nanopore Technologies R10.4.1 (R10). **B.** Mean per-read observed accuracy for WGS and WGA datasets from each species and sequencing platform. Observed read accuracy was calculated from read-to-reference alignments. **C.** Difference in mean per-read observed accuracy between paired WGA and WGS datasets. Positive values indicate lower mean accuracy in WGS than in the corresponding WGA dataset. **D.** Difference in mean per-read insertion, deletion, and substitution error rates between paired WGS and WGA datasets, calculated as WGS minus WGA. Positive values indicate higher error rates in WGS, whereas negative values indicate higher error rates in WGA. Differences are expressed in percentage points.

All nanopore datasets provided deep coverage and near-complete reference recovery. CS sequencing depth ranged from 184- to 884-fold, whereas R10 reads’ coverage exceeded 100-fold for every dataset except the Eco WGA sample, which reached 66-fold (**Fig. S1**). Across platforms and library types, more than 83.5% of primary mapped reads had a mapping quality of 60 (**Fig. S2**), and the aligned reads covered more than 99.9% of the corresponding reference (**Fig. S3**). These quality-control metrics minimized the likelihood that differences between WGS and WGA reads or platforms were attributable to insufficient sequencing depth, poor alignment, or incomplete genome representation.

### Native and amplified DNA exhibit distinct nanopore read-error profiles

Following quality control, we compared observed read accuracy between matched native WGS and WGA reads to assess methylation-associated differences at the whole-read level. Observed read accuracy was derived from read-to-reference alignments (**See Methods**).

WGA reads were generally more accurate than their matched WGS counterparts. For CS, WGA reads had higher mean observed accuracy in all six species, with WGS accuracy reduced by up to 0.298 percentage points relative to WGA (**Fig. 1B-C**). Four of the six R10 datasets showed the same direction of change, with the largest difference observed in Kpn, where WGS accuracy was approximately 0.577 percentage points lower than that of WGA (**Fig. 1C**). Additionally, median and mode values of observed read accuracy also showed similar results to the mean value (**Fig. S4**).

To identify the error types underlying these differences, we partitioned alignment errors into insertions, deletions, and substitutions. Across all CS datasets, WGS reads showed higher substitution and insertion rates but lower deletion rates than the corresponding WGA reads (**Fig. S5**). The WGS-associated increase in substitution rate ranged from 0.127 to 0.231 percentage points and consistently exceeded the corresponding increase in insertion rate, which ranged from 0.108 to 0.150 percentage points (**Fig. 1D**). Error profiles were more variable among the R10 datasets; nevertheless, substitutions were the largest positive component of the difference between WGS and WGA reads in Efa and Kpn, increasing by 0.337 and 0.385 percentage points, respectively (**Figs. 1D** and **S5**). In brief, the reduction in native-read accuracy was not attributable to a uniform increase across all error types; substitutions in the datasets showed the biggest differences between WGS and WGA reads.

### A2G and G2A substitutions form a recurrent nanopore error signature

Because substitutions accounted for a substantial component of the WGS-associated accuracy loss, we next decomposed substitution errors into the 12 possible classes. To account for differences in nucleotide composition, each class was quantified as the number of errors per callable reference base of the corresponding nucleotide without consideration of insertions and deletions (**see Methods**). Across all six species, A2G and G2A substitutions showed consistently elevated rates in both CS and R10 datasets (**Fig. 2A**). In CS reads, A2G and G2A error rates ranged from 0.15 to 0.66% and 0.23 to 0.52%, respectively; the corresponding ranges in R10 reads were 0.10-0.28% and 0.17-0.38% (**Fig. 2A**). The maximum enrichment of either class relative to the mean of the remaining ten classes was 33.9-fold for CS and 7.21-fold for R10.

**Figure 2.**
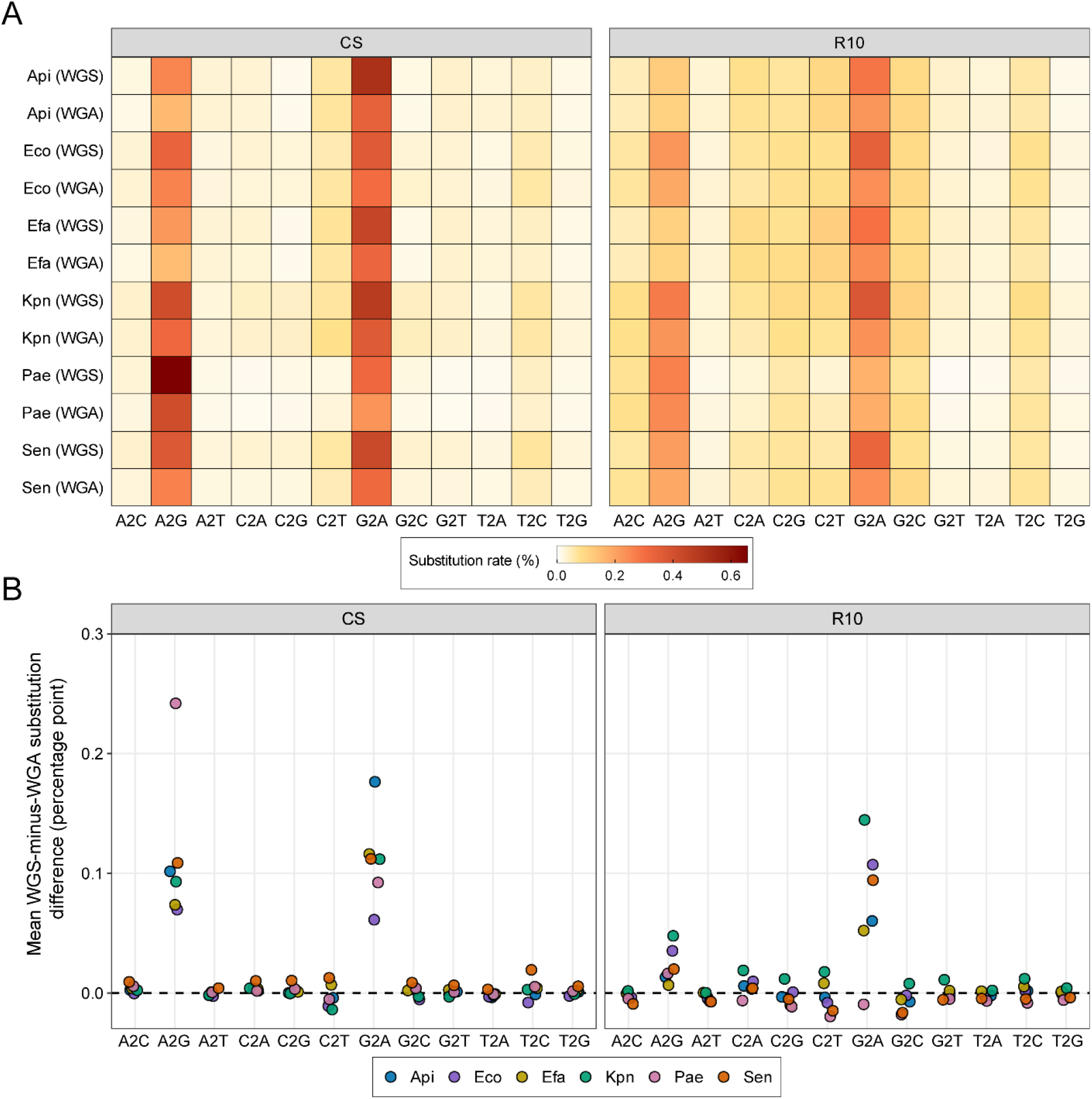
Adenine-to-guanine and guanine-to-adenine substitutions form a recurrent nanopore error signature. **A.** Substitution error rates for the 12 substitution types in paired WGS and WGA reads. For each substitution type, the error rate was calculated as the number of observed substitution events divided by the number of callable reference bases. **B.** Difference in substitution error rate between paired WGS and WGA reads for each of the 12 substitution types, calculated as WGS minus WGA. Each point represents one bacterial species. Positive values indicate a higher substitution error rate in native WGS reads, whereas negative values indicate a higher rate in the corresponding WGA reads. Differences are expressed in percentage points.

Row-wise z-score transformation of the substitution rates preserved the A2G/G2A-dominated pattern across species and DNA libraries (**Fig. S6**). The same signature was observed in data generated with the earlier CS configuration and ONT R9 platform, indicating that it was not restricted to the current sequencing configurations (**Fig. S7**). Re-basecalling the Efa WGS datasets with 12 ONT models spanning fast (FAST), high-accuracy (HAC), and super-accuracy (SUP) also retained the signal: G2A was the most strongly enriched class across all models, with A2G also elevated relative to most other substitution classes (**Fig. S8** and **Table S2**). By contrast, PacBio reads did not show a consistent cross-species A2G/G2A-dominated substitution spectrum (**Fig. S9**).

To determine whether methylation in native bacterial DNA contributed to this recurrent signature, we first calculated the paired WGS-minus-WGA difference for each substitution class. Differences for most of the other ten classes clustered around zero, whereas A2G and G2A showed the largest positive shifts, indicating higher substitution rates in native WGS reads (**Fig. 2B**). Across the six species, the mean differences in CS reads were 0.115 and 0.112 percentage points for A2G and G2A, respectively. In R10 reads, the corresponding differences were 0.023 and 0.075 percentage points, with the mean G2A effect exceeding the A2G effect by 0.052 percentage points (**Fig. 2B**).

Together, these analyses identify A2G and G2A enrichment as a recurrent substitution pattern in the tested nanopore datasets. This pattern remained detectable after whole-genome amplification, while native WGS reads showed an additional increase relative to matched WGA reads.

### Native DNA methylation is associated with localized A2G and G2A substitution errors

To determine whether the WGS-associated increase in substitutions was concentrated in methylated sequence contexts, we used PacBio data for independent methylation profiling and *de novo* motif identification. Eight distinct motifs were identified across ten species-motif combinations, with GATC detected in Eco, Kpn, and Sen (**Fig. 3A**). For each motif and its reverse complement, we defined a motif-centered unit encompassing the motif and 5 bp of flanking sequence on either side, which accounts for 4.18% of the whole chromosome. All remaining callable chromosomal positions were classified as non-motif regions (**Fig. 3A**). We then compared callable-base-normalized substitution rates between paired WGS and WGA datasets within these two genomic contexts (**See Methods**).

**Figure 3.**
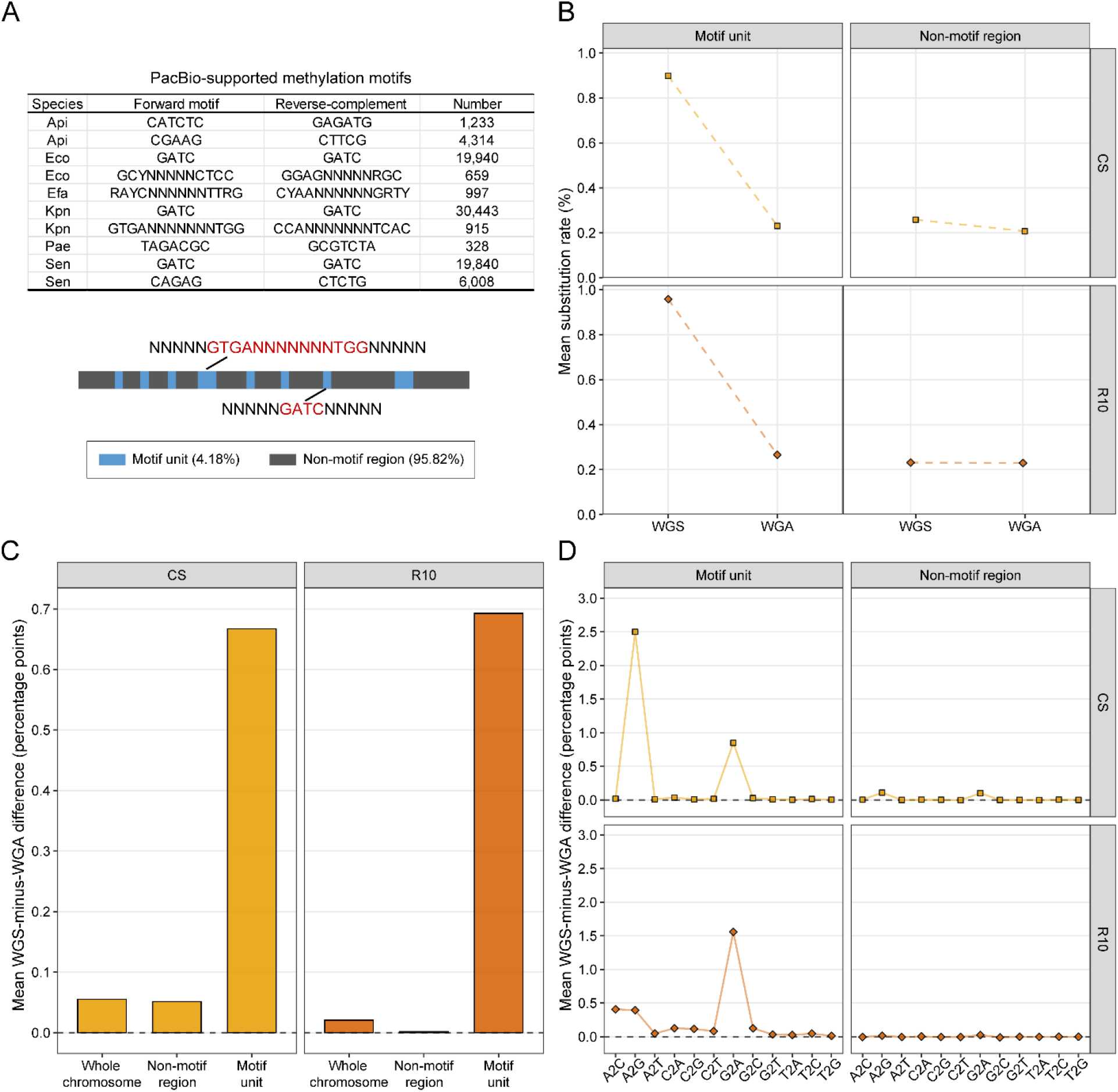
WGS-associated substitutions are concentrated around bacterial methylation motifs. **A.** Methylation motifs independently supported by PacBio data. The number indicates the total number of genomic occurrences of each motif, including occurrences of its reverse complement, in the corresponding chromosome. For the palindromic GATC motif, each genomic occurrence was counted only once because the forward and reverse-complement sequences are identical. For downstream analyses, each motif occurrence together with 5 bp of flanking sequence on either side was defined as a motif unit. All remaining chromosomal positions were classified as non-motif regions. **B**. Mean substitution rates in motif units and non-motif regions for WGS and WGA among six species. **C.** Mean difference in substitution rate between paired WGS and WGA datasets, calculated as WGS minus WGA, for the whole chromosome, non-motif regions, and motif units. **D.** Mean WGS-minus-WGA difference for each of the 12 substitution types within motif units and non-motif regions. Positive values indicate enrichment of the corresponding substitution type in WGS relative to WGA.

For both CS and R10, the WGS-associated increase in substitution rate was strongly concentrated within motif units. In CS, the mean substitution rate decreased from approximately 0.90% in WGS reads to 0.23% in WGA reads, corresponding to a WGS-minus-WGA difference of 0.668 percentage points (**Fig. 3B-C**). The corresponding R10 rates were approximately 0.96% and 0.27%, yielding a difference of 0.698 percentage points (**Fig. 3B-C**). By contrast, the WGS-minus-WGA difference in non-motif regions was only 0.051 percentage points for CS and approximately 0.002 percentage points for R10 (**Fig. 3B-C**). Relative to the whole-chromosome average, the motif-associated difference was approximately 12-fold greater in CS and 34-fold greater in R10 (**Fig. 3C**). These results indicate that the excess substitutions in native WGS reads were localized to methylation contexts rather than distributed uniformly across the chromosome.

We next examined which directional substitution types were elevated within these regions. Each type was normalized by the number of callable reference bases corresponding to its source nucleotide (**See Methods**). In CS, A2G showed the largest WGS-minus-WGA increase within motif units, reaching 2.502 percentage points, followed by G2A at 0.850 percentage points (**Fig. 3D**). In R10, G2A was the most strongly elevated class at 1.573 percentage points, whereas A2C and A2G showed similar secondary increases of 0.409 and 0.398 percentage points, respectively (**Fig. 3D**). The corresponding differences outside motif regions were substantially smaller, with no substitution types exceeding approximately 0.109 percentage points in CS or 0.024 percentage points in R10 (**Fig. 3D**). Thus, A2G and G2A accounted for the principal directional biases associated with methylated contexts, although R10 also exhibited a detectable A2C component.

The results of the WGS-minus-WGA difference varied considerably among individual motifs. In CS, the large aggregate A2G signal was influenced strongly by the Pae TAGACGC/GCGTCTA motif, at which the A2G difference reached approximately 12.149 percentage points (**Fig. S10**). At the same motif in R10 data, A2C showed the largest difference, at approximately 1.910 percentage points, followed by G2A at approximately 1.008 percentage points, whereas the A2G difference was smaller (**Fig. S11**). Across several other R10 motifs, G2A was the dominant substitution class (**Fig. S11**). These observations reveal motif- and platform-specific error spectra, indicating that the same methylated sequence context can be translated into different basecalling errors by different nanopore platforms (**Figs. S10** and **S11**).

To complement the motif-centered analysis, we asked whether the most pronounced WGS-associated substitution errors localized to methylated sequence contexts. Within each species-platform-substitution combination, we selected the top 1% of genomic sites ranked by the site-level WGS-minus-WGA A2G or G2A difference and compared their occurrence in PacBio-supported motif-centered regions with that of depth-matched control sites (**See Methods**). Of the 40 motif-platform-substitution comparisons, 39 remained significant after global multiple-testing correction (**Fig. S12**). Median enrichment was 4.70-fold and 3.71-fold for CS A2G and G2A sites, respectively, and 3.36-fold and 6.79-fold for the corresponding R10.4.1 sites. The strongest enrichment was observed for R10.4.1 G2A sites at the Eco GCYNNNNNCTCC 4mC-associated motif, which showed a 17.6-fold enrichment. Thus, genomic sites with the largest WGS-associated excess of A2G or G2A errors were preferentially concentrated in independently supported methylation contexts, although this enrichment establishes an association rather than a direct causal effect of methylation.

### Motif-associated substitutions produce high-fraction SNP-like discrepancies

Having established that the WGS-associated excess of substitutions was concentrated in methylation motif contexts, we next asked whether this signal was broadly distributed across motif occurrences or concentrated at a subset of loci. Within motif regions, we calculated the alternative-base fraction at each reference position and counted unique WGS-associated sites exceeding a series of thresholds (**See Methods**). As expected, the number of retained sites declined with increasing cutoff; nevertheless, high-fraction signals persisted. At a cutoff alternative-base fraction of 50%, 25 CS and 240 R10 sites remained across the six species (**Fig. 4A**).

**Figure 4.**
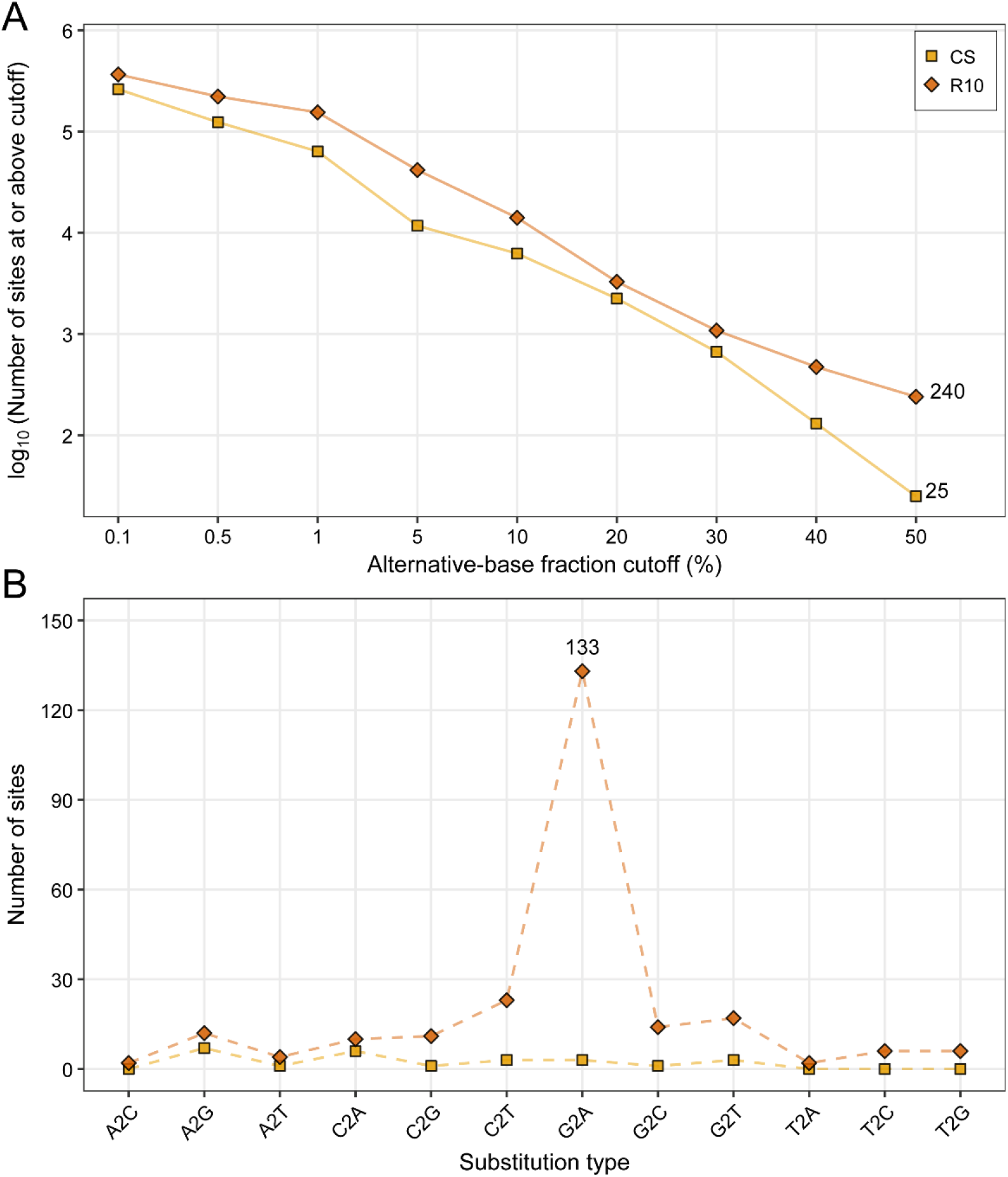
A subset of methylation-associated sites retains a high alternative-base fraction. **A.** Reverse cumulative site-count curves showing the number of motif-associated genomic positions with an alternative-base fraction greater than or equal to each cutoff. The number of sites retained for each cutoff was calculated from the number of sites with an alternative-base fraction greater than or equal to the cutoff in WGS reads but smaller than the cutoff in WGA reads. **B.** Distribution of substitution types among the sites retained at the 50% alternative-base fraction cutoff. For each site, the substitution type with the highest alternative-base fraction was defined as the representative substitution type.

In a haploid bacterial genome, an alternative-read fraction above 50% provides majority read support for alternative bases and may therefore resemble a single nucleotide polymorphism (SNP), although base fraction alone does not constitute a variant call. Among the retained CS sites, no single substitution type prevailed. By contrast, 133 of the 240 R10 sites (55.4%) were dominated by G2A substitutions (**Fig. 4B**).

### Methylation-associated strand-biased errors can confound ONT SNP calling

Motivated by the high alternative-base fractions observed at a subset of motif-associated loci and the strong G2A enrichment in R10, we next asked whether these errors could be retained as variants by the SNP-calling workflow (**See Methods**). After filtering with mapping quality (>=30), read depth (>=10), and variant QUAL (>=20), a total of 40 distinct PASS calls were observed across the R10 WGS, matched R10 WGA, and PacBio control callset (**Fig. 5A**). Of the 31 calls detected in WGS, 28 were unique to WGS; the remaining three were shared with WGA, and none overlapped the PacBio callset. These 28 WGS-specific calls were distributed across Api (n = 8), Eco (n = 10), Efa (n = 4), and Kpn (n = 6), with none detected in Pae or Sen (**Fig. 5B**). All 28 WGS-specific SNP calls retained after filtering were subsequently found to lie within the predefined methylation motif units. GATC accounted for the largest number of calls (n = 9) and encompassed six reference-oriented substitution classes. CATCTC accounted for a further eight calls, all of which were C2T substitutions (**Fig. 5C**).

**Figure 5.**
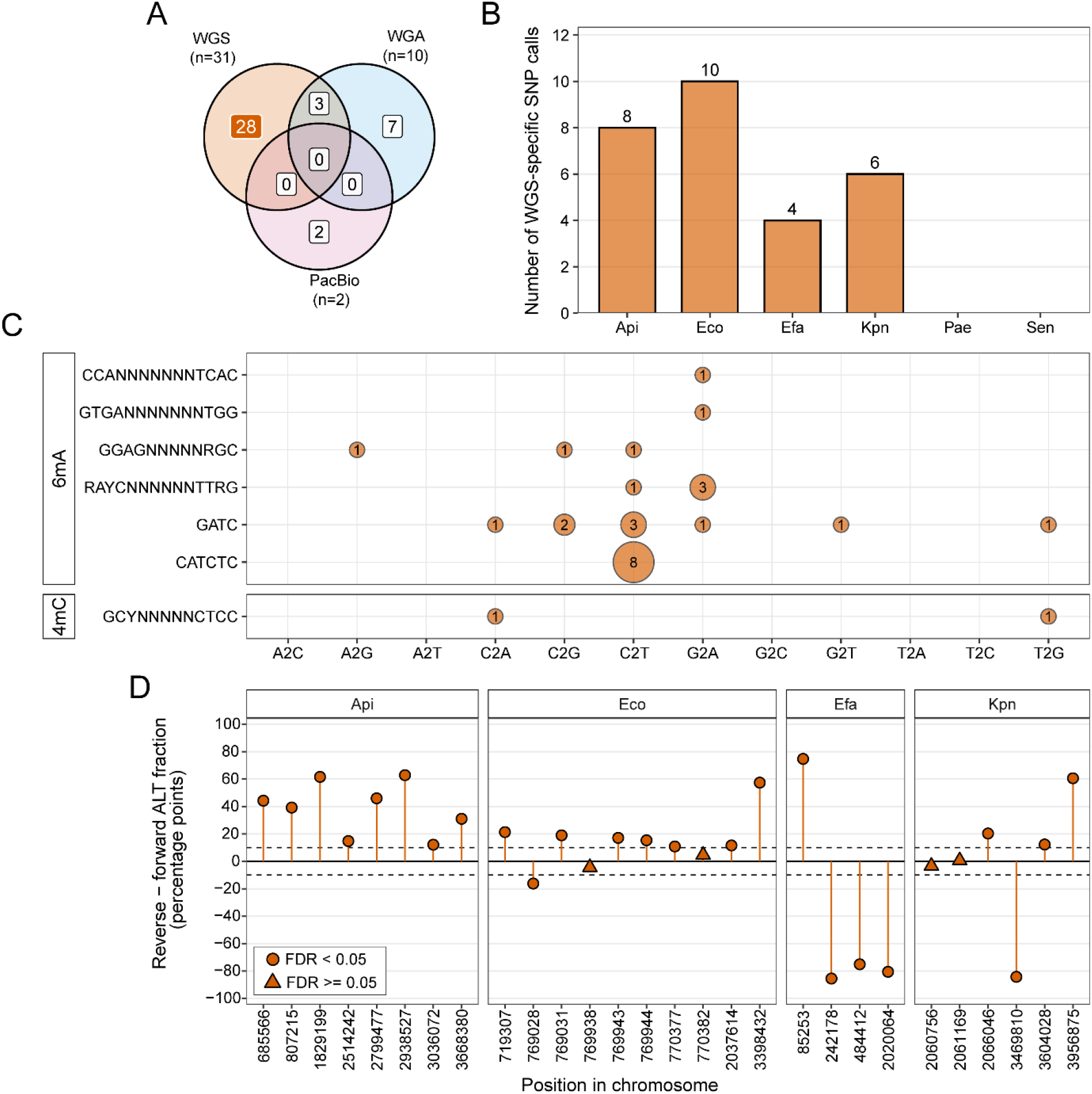
Methylation-associated strand-biased errors can survive R10 SNP-calling filters. **A.** Overlap of PASS SNP calls among R10 WGS, matched R10 WGA, and PacBio datasets. **B.** Distribution of R10 WGS-specific PASS SNP calls across the six bacterial species. A total of 28 WGS-specific calls were identified across Api, Eco, Efa, and Kpn. **C.** Relationship between methylation motifs, methylation types, and SNP types for the 28 WGS-specific PASS calls. All 28 calls were located within the methylation motif units. Circle size indicates the number of SNP calls for each motif-substitution combination. **D.** Strand bias of the 28 SNP calls, quantified from DP4 counts as the difference between reverse- and forward-strand alternative-allele fractions. Positive values indicate enrichment of the alternative allele among reverse-strand reads, whereas negative values indicate enrichment among forward-strand reads. Circles indicate calls with FDR < 0.05, and triangles indicate calls with FDR >= 0.05. Dashed lines indicate the predefined strand-bias effect-size threshold.

As methylation-associated nanopore errors can be strand-biased (Liu et al. 2024a), we quantified strand imbalance from the DP4 counts as the difference between reverse- and forward-strand alternative-allele fractions. Twenty-four of the 28 calls (85.7%) showed significant strand imbalance after false-discovery-rate correction, and 15 calls (53.6%) had an absolute strand-bias difference of at least 20 percentage points. All eight Api calls associated with the CATCTC motif showed greater alternative-allele support among reverse-strand reads, with six exceeding the 20-percentage-point threshold (**Fig. 5D**). An absolute strand-bias cutoff of 10 percentage points would flag 24 of the 28 methylation-associated calls, whereas a cutoff of 20 percentage points would flag 15 (**Fig. S13**). Thus, strand imbalance and proximity to independently supported methylation motifs provide complementary evidence for identifying putative methylation-associated SNP artifacts that survive the applied variant-calling filters.

## Discussion

Nanopore sequencing can directly interrogate native DNA and therefore preserves biological information such as DNA methylation, but the modifications can also complicate canonical basecalling (Chiou et al. 2023; Liu et al. 2024a; Thomas et al. 2025). In this study, using paired native WGS and methylation-depleted WGA libraries from six bacterial species, we characterized this effect from the whole-read level to individual SNP calls. WGA reads generally showed higher mean observed accuracy than their matched WGS reads, and substitutions explained much of this difference in the datasets with the largest accuracy loss (**Fig. 1**). At the nucleotide level, A2G and G2A substitutions formed a recurrent error signature across species, nanopore platforms, sequencing configurations, and basecalling models (**Figs. 2** and **S6-8**). Importantly, native WGS reads showed a further elevation of the A2G and G2A bias, and this increase was concentrated around PacBio-supported methylation motifs (**Fig. 3**). Among these loci, a subset of the alternative-base fraction exceeded 50%, and some signals subsequently survived an R10 SNP-calling workflow as WGS-specific PASS calls, most of which also exhibited significant strand imbalance (**Figs. 4** and **5**). In conclusion, we identify A2G and G2A as a shared nanopore-associated substitution bias and show that native bacterial DNA methylation further elevates this pre-existing bias at specific sequence contexts, in some cases producing SNP-like artifacts.

A key distinction emerging from our results is that methylation-associated substitutions occur on top of a broader and structured nanopore error spectrum. Nanopore substitution errors have long been recognized as non-random and have motivated the development of basecalling models trained using native methylated bacterial DNA (Delahaye and Nicolas 2021; Sanderson et al. 2024). In our study, A2G and G2A remained among the most enriched substitution types after amplification (**Fig. 2A**). These observations suggest that these two substitution types represent a shared nanopore-associated substitution bias rather than a methylation-specific phenomenon. Nevertheless, additional elevation of substitutions in native WGS relative to WGA indicates a second component associated with native DNA (**Fig. 2B**). This interpretation is consistent with previous studies showing that removal of DNA modifications through amplification can markedly reduce systematic nanopore errors (Chiou et al. 2023; Dabernig-Heinz et al. 2024).

The strongest evidence linking this additional component to methylation was its localization to independently supported methylation contexts (**Fig. 3**). The WGS-minus-WGA substitution difference was markedly elevated within PacBio-supported methylation motif units but was small in non-motif regions, and sites showing the largest WGS-associated A2G or G2A excess were preferentially enriched around these motifs (**Fig. 3C-D**). In an independent site-based analysis, genomic positions showing the largest WGS-associated A2G or G2A excess were also enriched around these motifs (**Fig. S12**). These observations extend previous reports that bacterial DNA modifications can generate errors at and around methylated positions (Liu et al. 2024a; Lohde et al. 2024; Thomas et al. 2025). The detailed error spectrum was platform-dependent. CS showed a stronger A2G component, whereas G2A was more prominent in R10 (**Fig. 3D**). Additionally, the same methylation motif, such as TAGACGC in Pae, can produce different dominant substitution types between platforms (**Figs. S10** and **S11**). This variation may arise because nanopore signals depend on the surrounding sequence, so a DNA modification can affect the signal of several nearby bases rather than only the modified base (Tourancheau et al. 2021). Differences in pore properties, signal processing, and model training may therefore determine how a similar modified sequence context is translated into a particular canonical-base error.

When we examined individual genomic positions, methylation-associated errors showed three clear features. First, only a subset of motif-associated positions developed high non-reference read fractions. The number of retained sites decreased rapidly as the alternative-base fraction increased, indicating substantial variation in error strength among individual positions (**Fig. 4**). Similar patterns have been reported in bacterial ONT data, where only a small proportion of methylated motif sites produced strong basecalling errors (Biggel et al. 2024; Thomas et al. 2025). Second, some of these error hotspots can reach a high alternative-base fraction and look like real SNPs. Unlike random sequencing errors, systematic errors can repeatedly occur at the same position in many reads. In our data, some motif-associated sites had an alternative-base fraction above 50%, and 28 WGS-specific calls passed the R10 SNP-calling workflow (**Fig. 5A-B**). All 28 calls were located within PacBio-supported methylation motif regions (**Fig. 5C**). This provides a direct link between methylation-associated read errors and SNP-like artifacts and is consistent with previous reports that methylation-related nanopore errors can affect genome assembly, cgMLST, and outbreak analysis (Biggel et al. 2024; Dabernig-Heinz et al. 2024; Lohde et al. 2024; Bogaerts et al. 2025). Third, many of these SNP-like errors showed strand bias. Most WGS-specific calls had different alternative-allele fractions between forward and reverse reads, consistent with previous reports of strand-specific methylation-associated errors (Liu et al. 2024a; Lohde et al. 2024). Therefore, these features, such as methylation motif proximity and strand bias, may be useful as complementary evidence for flagging suspicious SNP calls.

Several limitations should be considered in this study. First, WGA provides a practical methylation-depleted control but is not a methylation-only perturbation; amplification can alter other properties of the DNA library, and WGS-minus-WGA differences therefore cannot be attributed exclusively to methylation. The methylation interpretation in our analysis relies primarily on the evidence from independent PacBio motif profiling and motif-localized WGS-associated errors. Direct causality will require controlled experiments such as methyltransferase knockout and complementation or in vitro methylation of otherwise identical DNA templates. Second, the PacBio-supported motif set used here comprised 6mA- and 4mC-associated motifs. Important 5mC contexts, including CCWGG, were therefore not represented in the motif-related analyses, and our analysis may not capture the full contribution of bacterial DNA methylation to nanopore substitution errors. Third, the study included six bacterial strains and ten detected motifs, limiting separation of motif-specific effects from strain-specific genomic backgrounds. Finally, the variant-level analysis only used one R10 SNP-calling workflow. Because bacterial nanopore variant-calling performance can vary substantially among callers and basecalling configurations (Hall et al. 2024), the sensitivity of other callers, basecalling models, and sequencing depths to these errors remains to be established. Despite these limitations, our results indicate that bacterial DNA methylation should be considered not only as biological information retained by nanopore sequencing but also as a structured technical covariate affecting canonical basecalling and variant interpretation. Incorporating methylation context and strand-specific evidence into future nanopore analytical models may therefore improve bacterial variant reliability.

## Methods

### Biological resources and bacterial DNA collection

Six bacterial species, including *Acinetobacter pittii* (Api), *Escherichia coli* (Eco), *Klebsiella pneumoniae* (Kpn), *Pseudomonas aeruginosa* (Pae), *Salmonella enterica* (Sen), and *Enterococcus faecium* (Efa), were isolated and purified using methodologies described in prior publications (Zheng et al. 2007; Chen et al. 2017; Yang et al. 2019; Chen et al. 2020; Liu et al. 2022; Zhao et al. 2023). For each bacterial species, typical colonies were selected and cultured overnight in lysogeny broth (LB) at the appropriate growth temperature. Overnight cultures were then diluted 1:50 into fresh LB medium and incubated for an additional 4 h to reach log phase. After incubation, 2 mL of each culture was collected for genomic DNA extraction. The PureLink Genomic DNA Mini Kit from Invitrogen (USA) was used to extract the genomic DNA.

### CycloneSEQ library preparation and sequencing

Whole-genome amplification was performed as previously described (Ni et al. 2023). BGI personnel performed DNA library preparation and sequencing according to the manufacturer’s protocol. For library construction, approximately 500 ng of each sample was subjected to end repair by adding End Repair Buffer and End Repair Enzyme, followed by incubation at 20 degrees Celsius for 10 minutes and 65 degrees Celsius for 10 minutes, then held at 4 degrees Celsius. The mixtures were subsequently purified using 1.0× DNA Clean Beads and eluted in nuclease-free water. The end-repaired samples were then ligated with barcodes using Ligation Barcode, 4× Ligation Buffer, and DNA Ligase, with incubation at 25 degrees Celsius for 30 minutes. The ligation products were purified with 0.4× DNA Clean Beads and recovered in nuclease-free water. Next, barcoded samples were combined with Barcode Sequencing Adaptors, 4× Ligation Buffer, and DNA Ligase, and incubated again at 25 degrees Celsius for 30 minutes. A second purification step was performed using 0.4× DNA Clean Beads. The products were then treated with the long-fragment wash buffer to remove short fragments and finally eluted in the elution buffer.

The constructed libraries were quantified with a Qubit fluorometer and sequenced on the CycloneSEQ platform following the standard procedure. Two different combinations of library preparation kits and basecallers were used: the initial version involved the CycloneSEQ 24 Barcode Library Prep Set (H940-000018) with basecaller v1.1.0.9, while the updated version used the CycloneSEQ 24 Barcode Library Prep Set (H940-000081) with basecaller v1.3.0.23. All sequencing was carried out on the G100-ER platform.

### ONT read basecalling

Public ONT reads in FASTQ format were basecalled using Guppy (V6.4.6) with the dna_r10.4.1_e8.2_400bps_sup.cfg model for R10.4.1 and dna_r9.4.1_450bps_sup.cfg for R9.4.1 sequencing reads (Liu et al. 2024a; Ye et al. 2024). The Efa isolate was additionally sequenced at a sampling frequency of 5 kHz to provide input for the 5-kHz models. Efa raw signal datasets were re-basecalled using Dorado (V0.9.1) with twelve ONT models covering FAST, HAC, and SUP accuracy modes (**Table S2**).

### Nanopore read processing and read-accuracy-related analysis

For all the nanopore data, reads with a length of less than 200 bp were filtered out. To remove potential contamination, the reads that were not aligned to the related reference were also filtered out. The filtered reads were regarded as clean data for the downstream analysis.

To calculate the genome covered rate at the chromosome level for each sample, the clean reads were aligned to the related reference using Minimap2 (V2.17-r941) (Li 2021) with the arguments “-ax map-ont”, and the generated BAM files were then profiled by the subfunction named “coverage” from SAMtools (V1.18) (Danecek et al. 2021). To calculate the observed metrics, including observed read accuracy and mismatch rate, the “observe” subfunction of Giraffe (V0.2.3) (Liu et al. 2024b) was used with the BAM files as inputs. During the calculation of Giraffe, the number of insertions and deletions was included for analysis as follows:

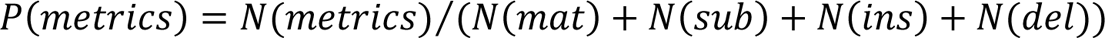

Here, the metrics can be mat, sub, ins, or del. N(*mat*), N(*sub*), N(*ins*), and N(*del*) are the number of matched bases, substitutions, insertions, and deletions.

### Callable-base-normalized substitution rate calculation

To quantify the percentage of substitution type rates independently of insertions and deletions, the subfunction of “mpileup” in SAMtools (V1.18) (Danecek et al. 2021) was utilized with the arguments of “--min-MQ 30 --min-BQ 0 --no-output-ins --no-output-ins --no-output-del --no-output-del --no-output-ends”. The argument “--min-MQ 30” was used to filter the aligned reads with mapping quality (mapQ) smaller than 30. The arguments “--no-out-del”, “--no-output-ins”, and “--no-output-ends” were used only to output the substitutions and matches. The default value of the “--min-BQ” argument is 13, which will skip the aligned base with an estimated quality (Q value) smaller than 13. We set the argument to 0 to keep all the aligned bases for mismatch number counting.

Substitutions were classified in the native orientation of each read. At a genomic position with a reference base, forward-aligned reads were evaluated relative to the reference base, whereas reverse-aligned reads were evaluated relative to its complementary base. Consequently, each reference position could contribute to six substitution types. For example, a genomic reference C position contributed C2A, C2G, and C2T from forward reads, and G2A, G2C, and G2T from reverse reads. The site-level fractions for C2A and G2A were calculated as follows:

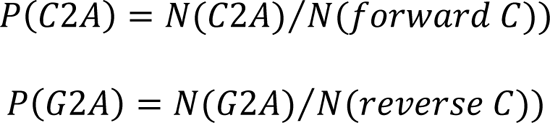

Here, N (C2A) denotes the number of forward-aligned reads with an observed A at the reference C position. N (G2A) denotes the number of reverse-aligned reads with an observed T at the same position in the alignment, corresponding to a G2A substitution in the native read orientation. N (forward C) and N (reverse C) denote the total number of callable read observations at that position from forward- and reverse-aligned reads, respectively, including both reference-matching and alternative bases.

For each genomic region, substitution counts were pooled across positions and alignment orientations and divided by the corresponding total number of callable read observations. For example, the regional C2A substitution rate was calculated as:

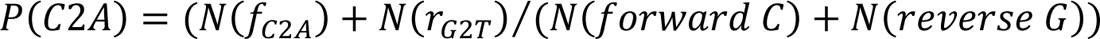

Here, N(f(C2A)) counts C2A in forward-aligned reads, whereas N(r(G2T)) counts G2T in reverse-aligned reads, with both labels expressed relative to the reference orientation. Both represent C2A substitutions in the native read orientation. The denominator comprises callable observations from forward-aligned reads at reference C positions and reverse-aligned reads at reference G positions. The same approach was applied to the other substitution types.

At each genomic position, the alternative-base fraction was calculated by combining callable observations from both alignment orientations:

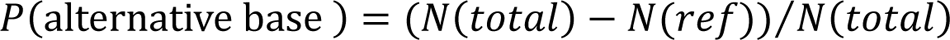

Here, N (total) denotes the total number of callable read observations at that position, and N (ref) denotes the number matching the reference base.

For regional WGS-minus-WGA comparisons, analyses were restricted to chromosomal positions with at least 10 callable read observations in each of the paired WGS and WGA datasets.

### Row-wise z-score normalization of substitution rates

To compare patterns across the 12 substitution types, we calculated z-scores separately for each dataset (one heatmap row). Each dataset represented a combination of species, sequencing platform, and DNA library type (WGS or WGA). Callable-base-normalized substitution rates were used without log transformation or adding pseudocounts. For substitution type (j) in dataset (i), the z-score was calculated as:

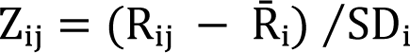

Here, R_ij_ is the callable-base-normalized substitution rate for type (j), and R^-^_i_ and SD_i_ are the mean and sample standard deviation of the 12 substitution rates in dataset (i).

Positive z-scores indicate rates above the dataset mean, whereas negative z-scores indicate rates below the mean. The z-score heatmap therefore shows which substitution types have relatively high or low rates within each dataset. Absolute differences between datasets were assessed using the original callable-base-normalized substitution rates. For PacBio data, the same normalization was performed separately for each species using its 12 substitution rates.

### Methylation motif detection through PacBio data

PacBio HiFi sequencing data were used to independently characterize DNA methylation patterns in the six bacterial isolates. Raw unaligned HiFi BAM files containing polymerase kinetic information were processed using SMRT Link (V26.1.0). The resulting reads were aligned to the corresponding chromosome reference using pbmm2 with the CCS preset, followed by BAM indexing. Plasmid sequences were excluded from this analysis. DNA modifications were identified using ipdSummary with the SP3-C3 kinetic model. Both 6mA and 4mC were evaluated using a significance threshold of P < 0.001, and the methylated-read fraction was estimated at each genomic position. Methylation motifs were inferred from the modification calls using pbmotifmaker with a minimum modification score of 35. Motifs with a methylated fraction greater than 99% were considered supported.

### Enrichment analysis of top 1% A2G and G2A error sites

We tested whether methylation motifs were enriched around sites with large WGS-minus-WGA substitution differences, separately for A2G and G2A substitutions. For each species and platform (CS or R10), WGS and WGA pileups were compared at matching chromosome positions and native-read orientations. A site was callable when the corresponding strand had at least 10 A/C/G/T observations in both datasets. The substitution-rate difference, expressed in percentage points, was calculated as:

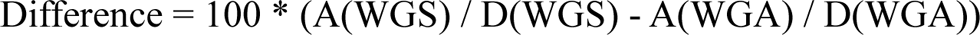

Here, (A) is the number of reads supporting the specified alternative base, and (D) is the strand-specific callable depth.

Within each species, platform, and substitution type, sites were ranked from highest to lowest WGS-minus-WGA difference. The target number of selected sites was 1% of all callable sites, rounded up, and only sites with positive differences were eligible. For each selected site, we extracted a 31-bp sequence in the native-read orientation, comprising the focal base and 15 bp on each side. Windows extending beyond chromosome boundaries or containing non-ACGT bases were excluded. An equal number of non-selected callable sites were sampled as controls from the same species, platform, and substitution type. Controls were matched by the lower of the WGS and WGA callable depths, grouped into five categories: 10-19, 20-39, 40-79, 80-159, and >= 160 reads. Enrichment of the ten predefined methylation-motif families was tested using SEA in MEME Suite (V5.5.7) (Bailey et al. 2015), separately for each species, platform, and substitution type. Enrichment ratios compared motif-hit frequencies between selected and control sequences. P-values from all 40 predefined motif tests were adjusted together using the Benjamini-Hochberg procedure. Adjusted P-values below 0.05 were considered significant. For complementary de novo motif discovery, sequences were pooled by platform and substitution type, with each species contributing up to 1,000 selected and 1,000 control sequences. STREME was used to identify up to five motifs, each 4-15 bp long. TOMTOM compared these motifs with the predefined motif database using a minimum overlap of four bases and a match threshold of 0.1.

### SNP calling and identification of WGS-specific calls

Variants were called separately from chromosome-aligned WGS, WGA, and PacBio BAM files using BCFtools (V1.23.1) (Danecek et al. 2021) and SAMtools/HTSlib (V1.23). BCFtools mpileup was run against the corresponding species chromosome reference with minimum mapping and base qualities of 30 and 0, respectively, and a maximum depth of 1,000,000. Unmapped, secondary, quality-control-failed, duplicate, and supplementary alignments were excluded. Read-group distinctions were ignored, and indels were skipped. Total read depth (DP) and total, forward-strand, and reverse-strand allelic depths (AD, ADF, and ADR) were retained. The BCFtools multiallelic caller was run under a haploid model, retaining only variant sites. Calls were normalized against the reference, multiallelic records were split, and only single-nucleotide substitutions were retained. The primary PASS criteria were QUAL >= 20 and read depth >= 10.

Before comparing call sets, we removed calls potentially caused by plasmid repeat reads mapping to the chromosome. SNP-centred sequence windows were searched against the polished chromosome and available plasmid sequences for matching repeat copies outside the original locus. Moderate matches required >= 80% sequence identity and >= 60% query coverage.

Calls were compared by species, chromosome, position, reference base, and alternative base. WGS-specific SNP calls were defined as repeat-filtered PASS alleles detected in WGS but absent as the same allele in both matched WGA and PacBio callsets. After these calls had been selected, their genomic positions were compared with the methylation motif units, comprising each motif and 5 bp of flanking sequence on either side.

Strand imbalance was evaluated for each WGS-specific PASS SNP using the forward- and reverse-strand alternative-read counts from the DP4 field. The alternative-allele fraction was calculated separately for forward and reverse reads as the alternative-read count divided by the total callable read count on that strand. Strand imbalance was defined as the reverse-strand alternative-allele fraction minus the forward-strand alternative-allele fraction. Statistical significance was assessed using a two-sided Fisher’s exact test on the forward/reverse reference- and alternative-read counts, and P values were adjusted across the 28 methylation-associated calls using the Benjamini-Hochberg procedure. Absolute strand-bias differences of 10 and 20 percentage points were used as descriptive thresholds; strand bias was not used as a PASS filtering criterion.

### Use of AI-assisted tools

OpenAI Codex and ChatGPT were used to assist with developing and refining scripts for data analysis and visualization. The authors reviewed the resulting code and checked the analytical outputs for accuracy. The authors take full responsibility for the analyses and their interpretation.

## Data access

All sequenced CycloneSEQ and PacBio datasets generated in this study have been deposited in the NCBI Sequence Read Archive under BioProject ID PRJNA980403. The related ONT reads are also available in the same BioProject.

All analysis outputs and corresponding genome assemblies have been uploaded to Figshare (https://figshare.com/projects/_b_Nanopore_bias_project_b_/281035).

## Acknowledgment

We thank Professor Chen Sheng (Hong Kong Polytechnic University) for providing bacterial DNA samples and Dr. Ye Lianwei (City University of Hong Kong) and Dr. Sun Yuanyang (Hong Kong Polytechnic University) for bacterial culture and sample preparation. We are also grateful to the staff of BGI Tech for conducting CycloneSEQ sequencing. ChatGPT (OpenAI) was used to assist with improving the wording, clarity, and organization of the manuscript. The authors reviewed and revised the AI-assisted text and took full responsibility for the final content.

This work was supported by the Early Career Scheme (CityU 21100521) and General Research Fund (11105524) from the Research Grants Council of the Hong Kong Special Administrative Region, China; the Hong Kong Health and Medical Research Fund (project number 08194126); and new Research Initiatives support from City University of Hong Kong (project number 9610497) to R.L; and was supported by the City University of Hong Kong (project number 9683001) to M.Y.

## Contribution

**Xudong Liu:** Conceptualization, methodology, validation, formal analysis, investigation, data curation, writing (original draft, review, and editing). **Qiutao Ding:** Methodology, data curation, and writing (review and editing). **Yanwen Shao:** Methodology, data curation, and writing (review and editing). **Zhihao Guo**: Methodology, data curation, and writing (review and editing). **Ying Ni**: Writing (review and editing). **Lu Fan**: Writing (review and editing). **Yi Yang**: Writing (review and editing). **Kaixin Chen**: Writing (review and editing). **Mengsu Yang**: Writing (review and editing), supervision, and funding acquisition. **Runsheng Li**: Conceptualization, methodology, validation, investigation, writing (original draft, review, and editing), supervision, and funding acquisition.

