## Supplementary materials for "Bacterial DNA methylation amplifies the shared A-to-G and G-to-A substitution bias in nanopore sequencing"

### Supplementary table

**Table S1. Information on the genome reference.**

| ID | Species | Type | Genebank | Size (bp) | GC content (%) |
|---|---|---|---|---|---|
| Api | *Acinetobacter pittii* | Gram-negative | CP127321.1 | 3,896,718 | 38.95 |
| Eco | *Escherichia coli* | Gram-negative | CP127324.1 | 4,868,913 | 50.55 |
| Kpn | *Klebsiella pneumoniae* | Gram-negative | CP127340.1 | 5,406,825 | 57.5 |
| Pae | *Pseudomonas aeruginosa* | Gram-negative | CP127342.1 | 6,410,639 | 66.4 |
| Sen | *Salmonella enterica* | Gram-negative | CP127343.1 | 5,019,577 | 52.17 |
| Efa | *Enterococcus faecium* | Gram-positive | CP127337.1 | 2,785,439 | 38.03 |

**Table S2. Information on the nanopore basecalling models used.**

| ID | Name | Platform |
|---|---|---|
| 5.2_fast | dna_r10.4.1_e8.2_400bps_fast@v5.2.0 | R10 |
| 5.2_hac | dna_r10.4.1_e8.2_400bps_hac@v5.2.0 | R10 |
| 5.2_sup | dna_r10.4.1_e8.2_400bps_sup@v5.2.0 | R10 |
| 5khz_fast | dna_r10.4.1_e8.2_400bps_fast@v5.0.0 | R10 |
| 5khz_hac | dna_r10.4.1_e8.2_400bps_hac@v5.0.0 | R10 |
| 5khz_sup | dna_r10.4.1_e8.2_400bps_sup@v5.0.0 | R10 |
| 4khz_fast | dna_r10.4.1_e8.2_400bps_fast@v4.1.0 | R10 |
| 4khz_hac | dna_r10.4.1_e8.2_400bps_hac@v4.1.0 | R10 |
| 4khz_sup | dna_r10.4.1_e8.2_400bps_sup@v4.1.0 | R10 |
| R941_sup | dna_r9.4.1_e8_sup@v3.6 | R9 |
| R941_fast | dna_r9.4.1_e8_fast@v3.4 | R9 |
| R941_hac | dna_r9.4.1_e8_hac@v3.3 | R9 |

#### Supplementary figures

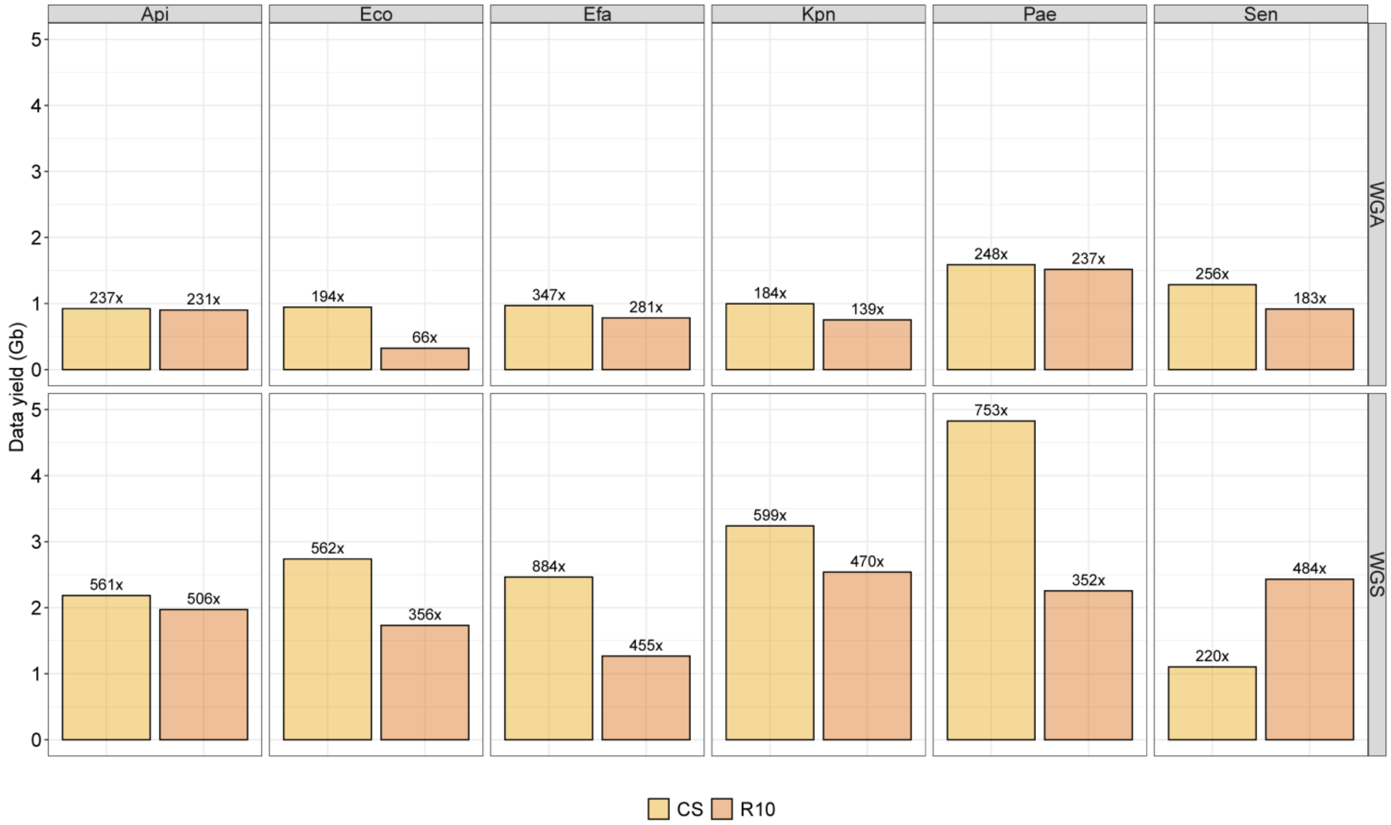


**Figure S1.** Clean-data yield for CycloneSEQ (CS) and Oxford Nanopore Technologies R10.4.1 (R10) datasets across the six bacterial species. Values above the bars indicate sequencing depth relative to the corresponding reference chromosome, expressed as fold coverage. WGS, whole-genome sequencing; WGA, whole-genome amplification.

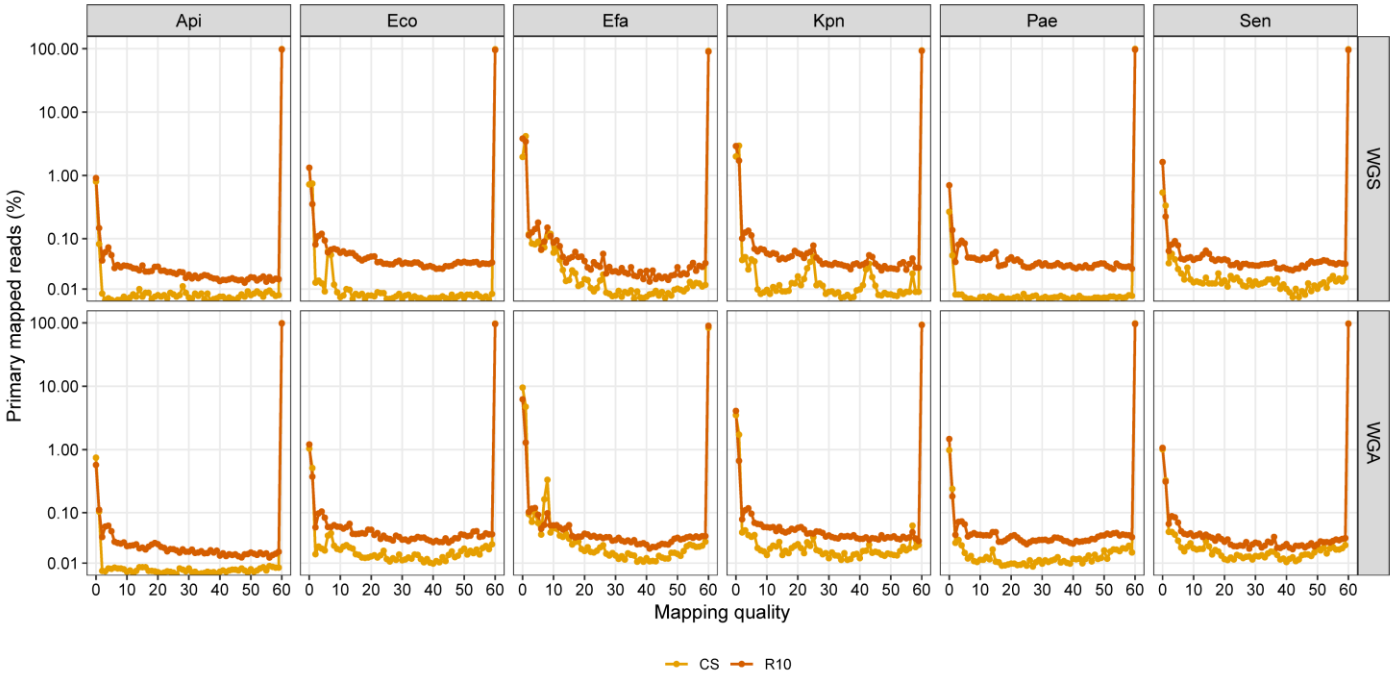


**Figure S2.** Distribution of mapping quality among primary mapped reads. The y-axis represents the percentage of primary mapped reads at each mapping-quality score and is shown on a logarithmic scale.

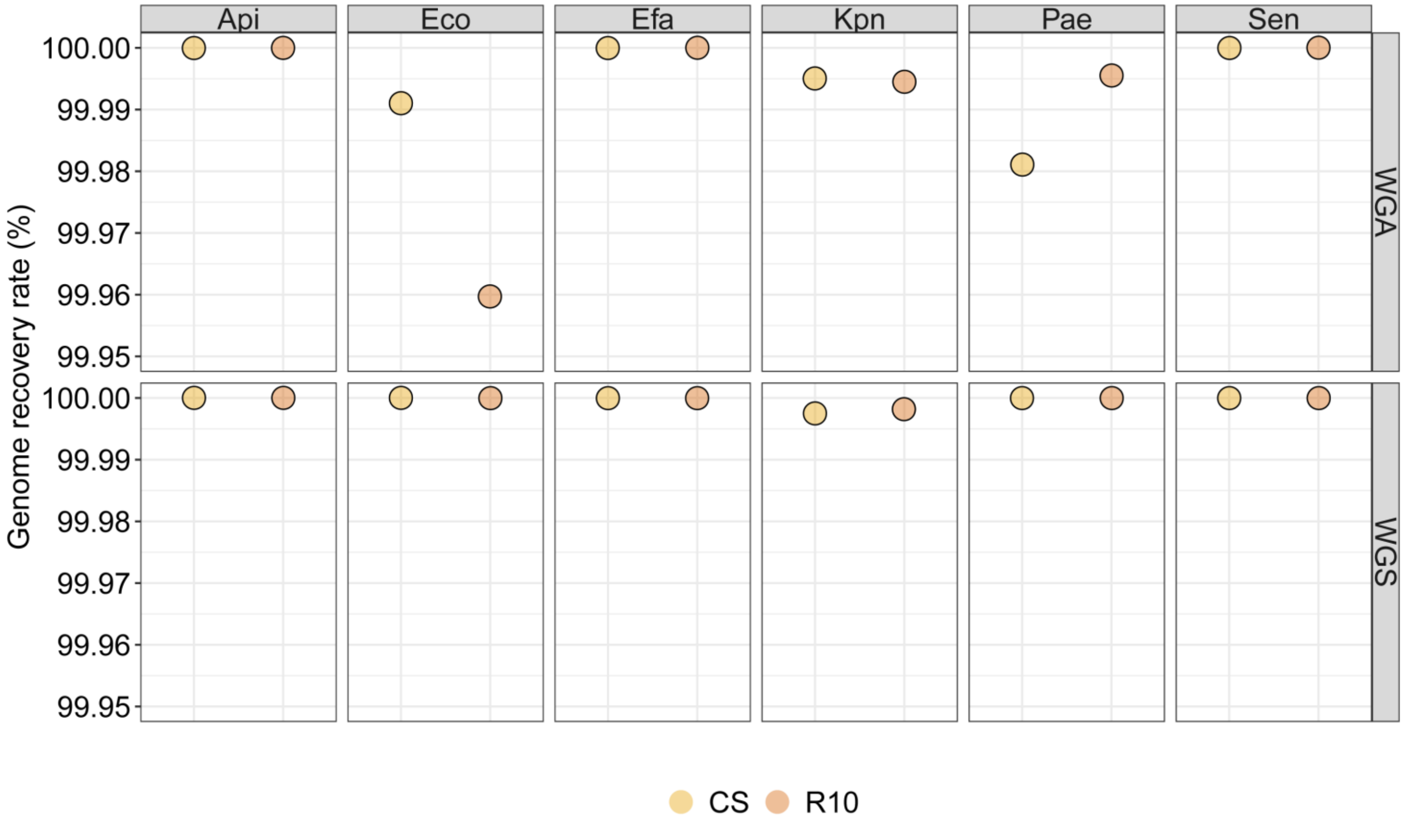


**Figure S3.** Chromosome-level genome recovery rates for CS and R10 datasets across the six bacterial species. Genome recovery represents the proportion of the corresponding reference chromosome covered by mapped reads.

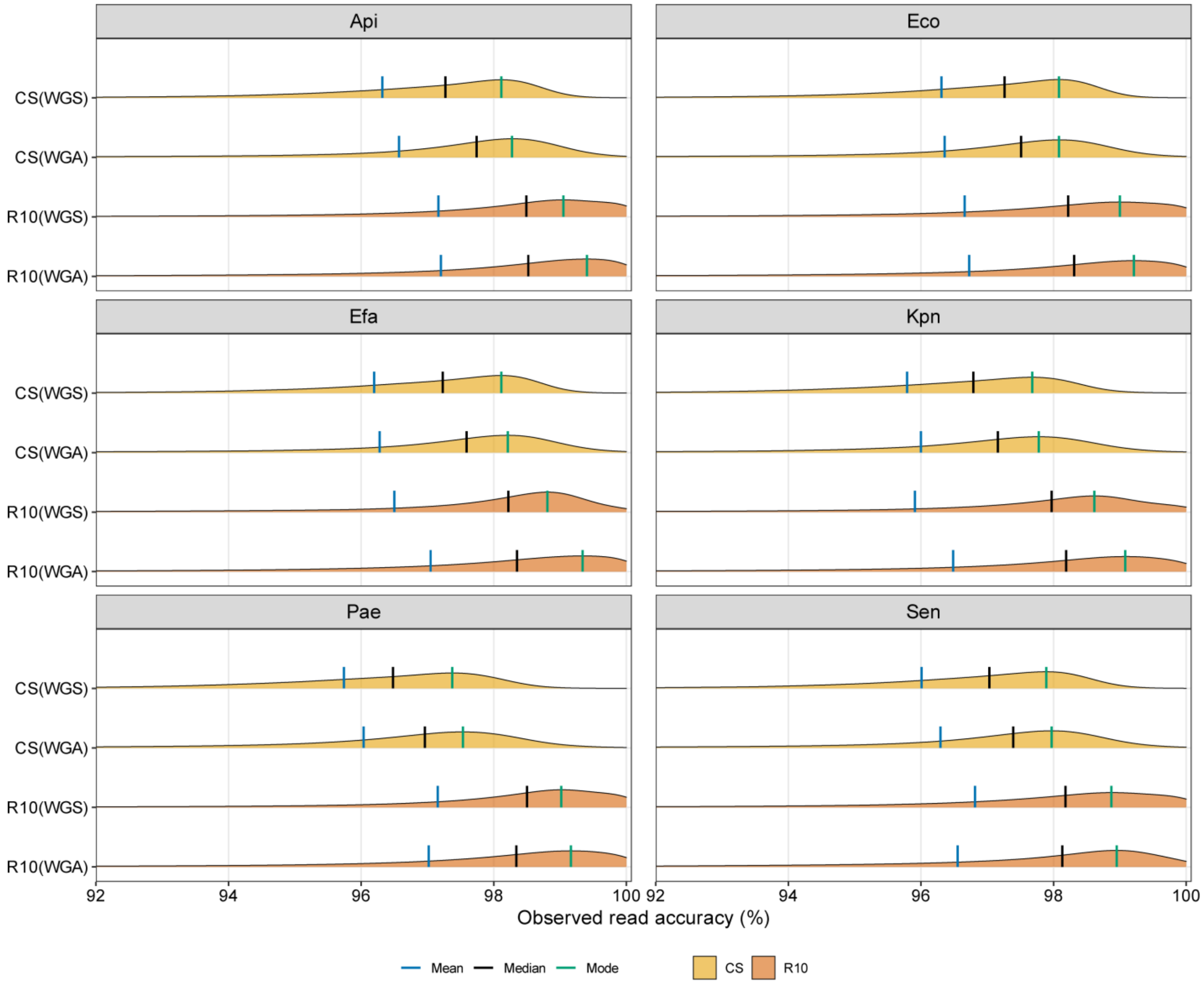


**Figure S4.** Per-read observed accuracy distributions for paired WGS and WGA datasets. Vertical lines indicate the mean, median, and mode of each distribution.

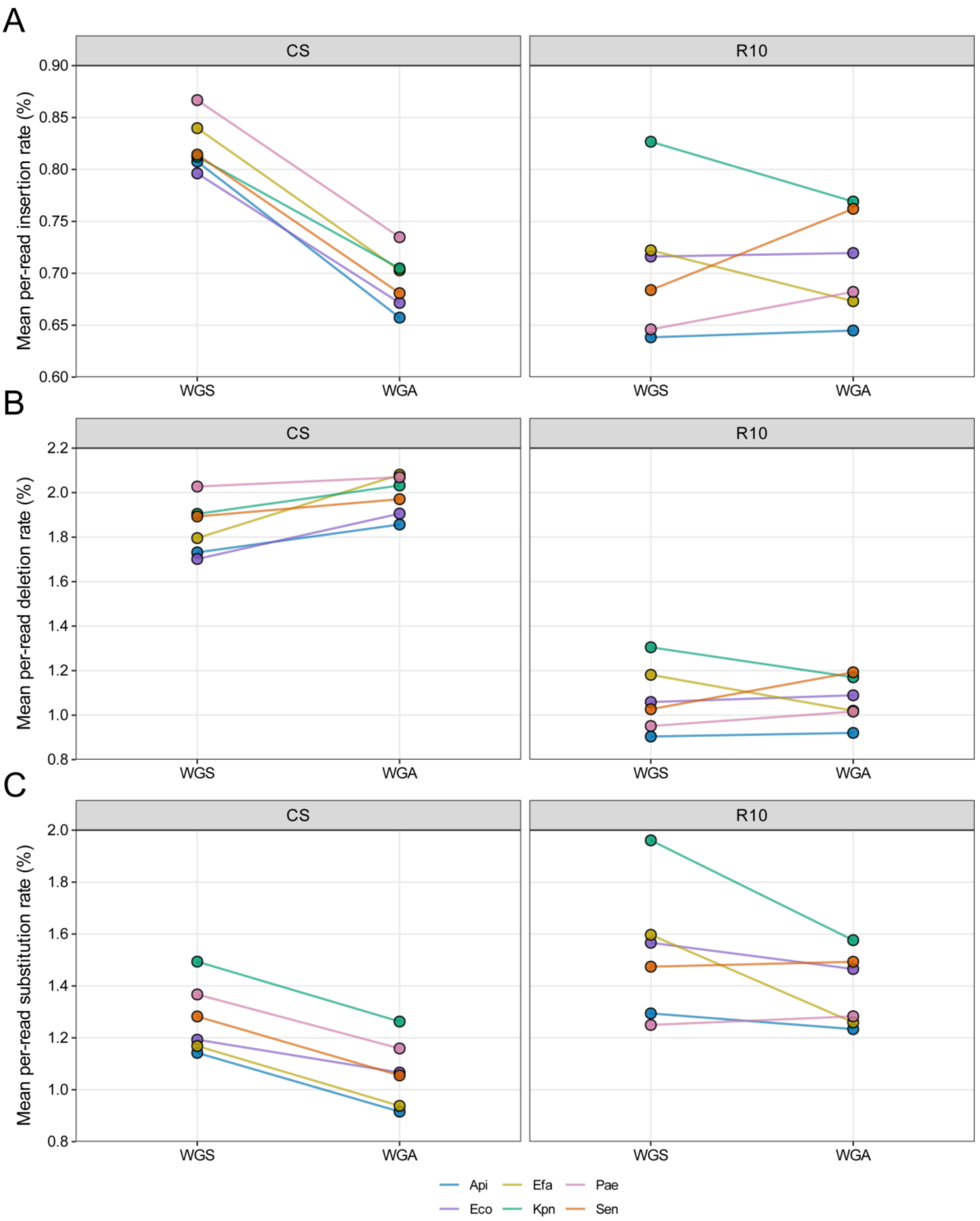


**Figure S5.** Mean per-read **A.** insertion, **B.** deletion, and **C.** substitution error rates for CS and R10 datasets. Lines connect paired WGS and WGA datasets from the same bacterial species. Each point represents the mean per-read error rate.

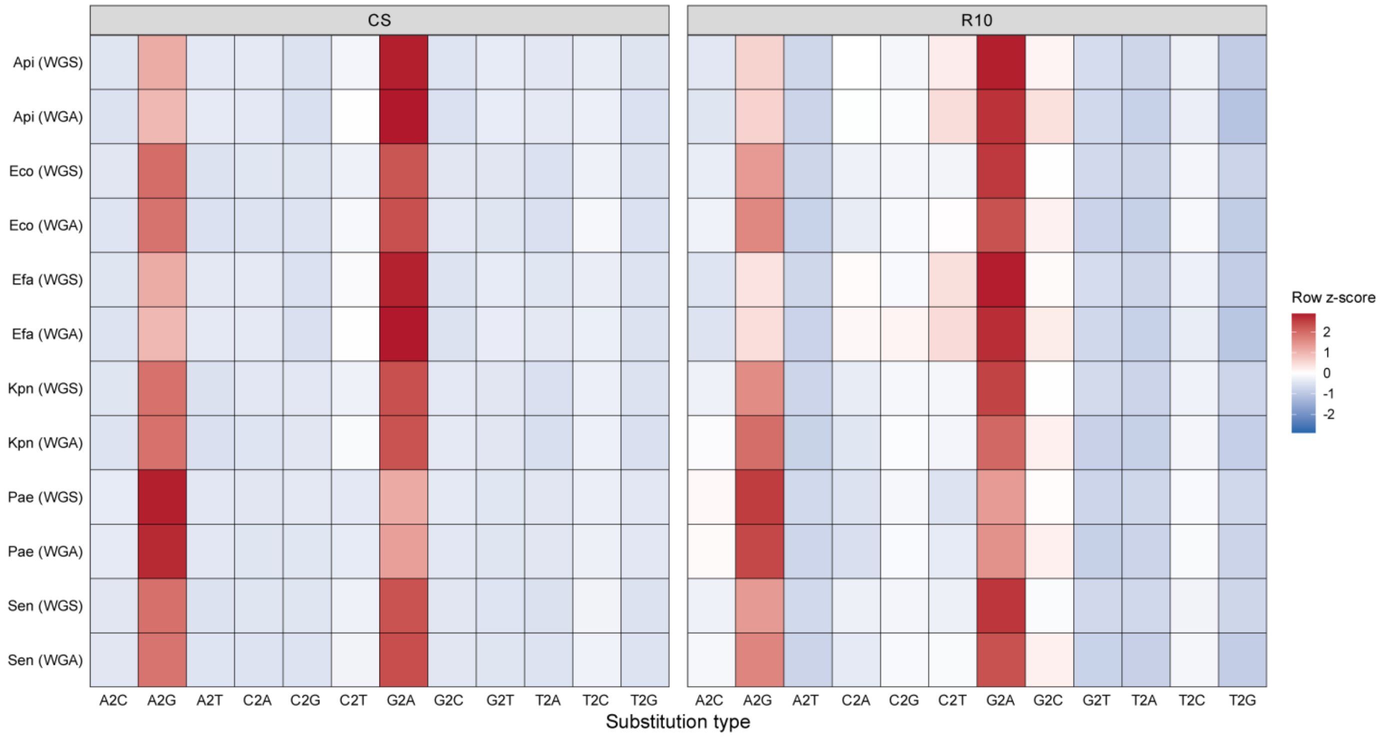


**Figure S6.** Row-wise z-score heatmaps of the 12 substitution types for CS and R10 datasets. For each dataset, substitution rates were transformed to z-scores across the 12 substitution types; positive values therefore indicate substitution types elevated relative to the other classes within the same dataset.

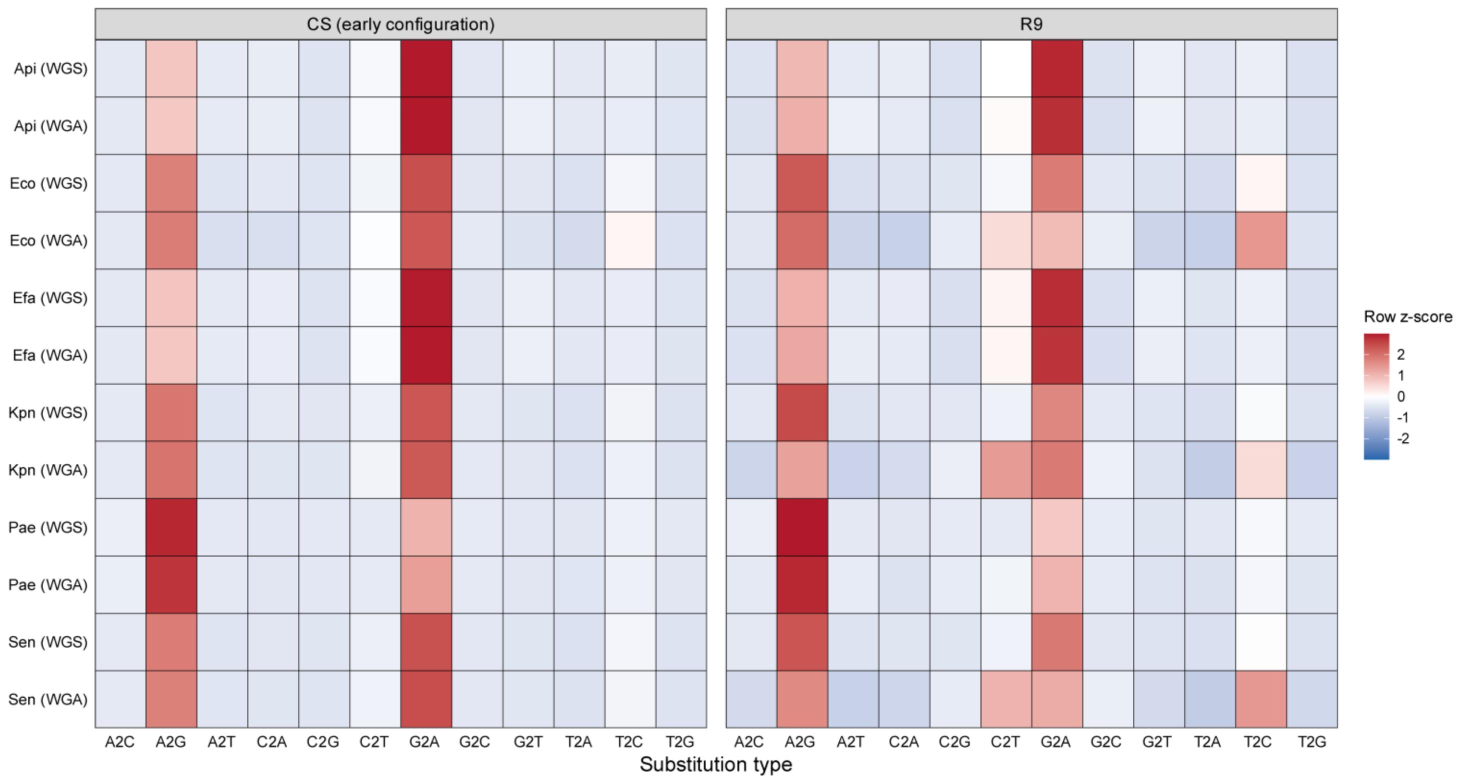


**Figure S7.** Row-wise z-score heatmaps of the 12 substitution types in paired WGS and WGA datasets generated using the earlier CS configuration and Oxford Nanopore Technologies R9.4.1 (R9).

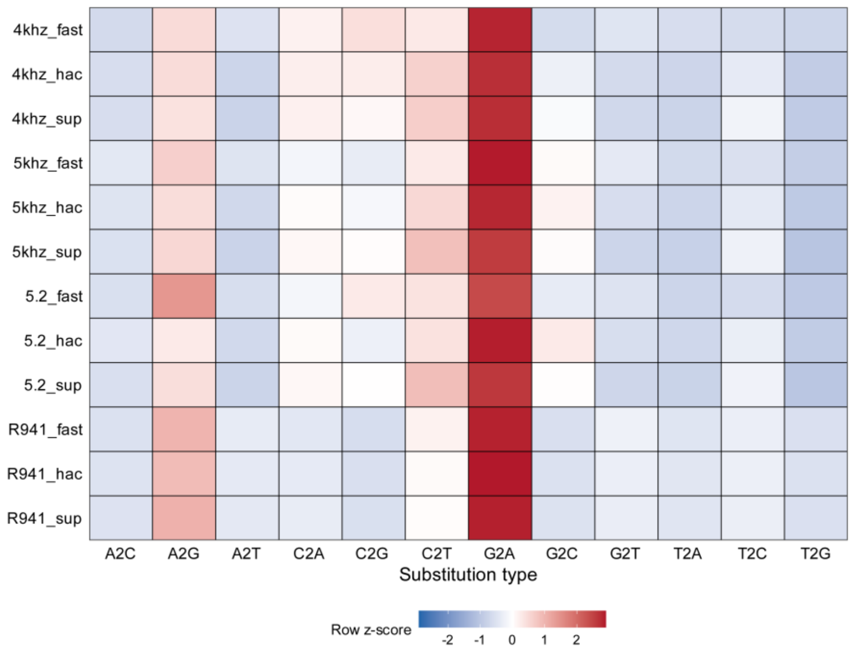


**Figure S8.** Row-wise z-score profiles of the 12 substitution types for ONT Efa WGS reads re-basecalled using 12 basecalling models spanning fast (FAST), high-accuracy (HAC), and super-accuracy (SUP) configurations from R10 and R9. Each row represents one basecalling model, and z-scores were calculated across the 12 substitution types within that model.

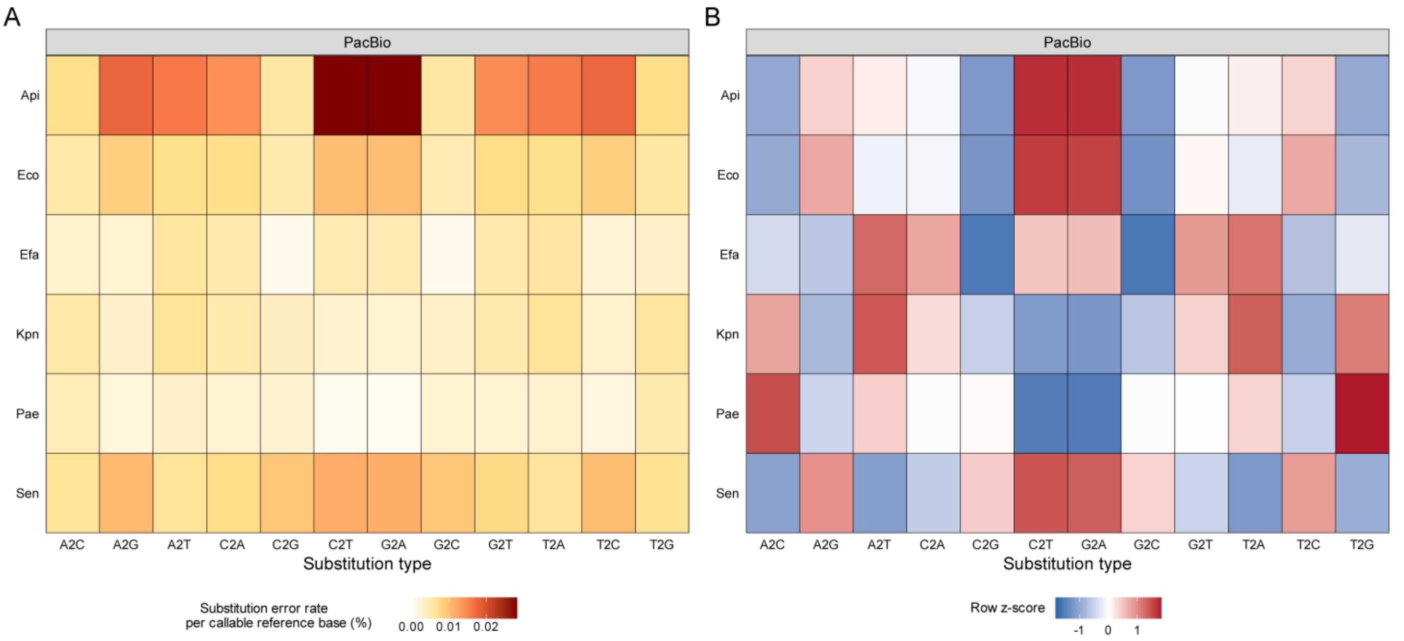


**Figure S9. A.** Substitution rates for the 12 substitution types in PacBio reads. **B.** Row-wise z-score transformation of the substitution rates shown in A, calculated across the 12 substitution types for each species.

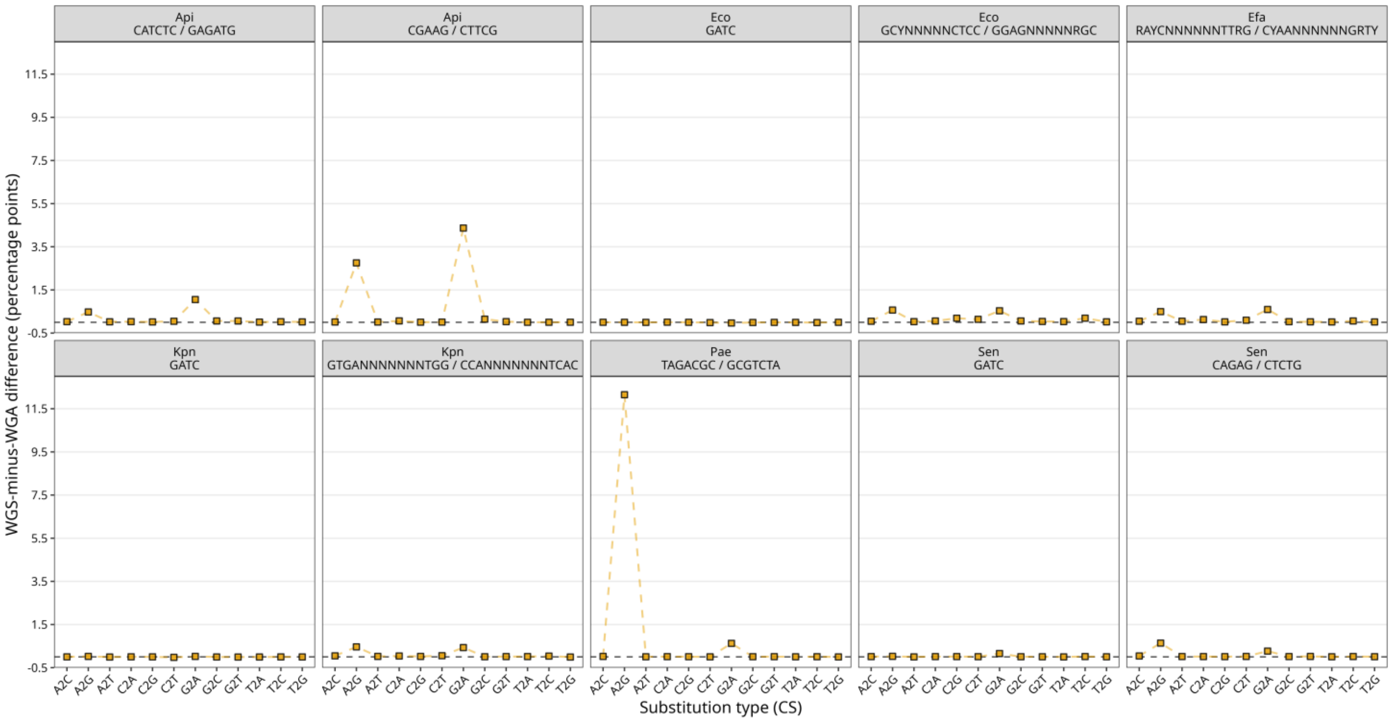


**Figure S10.** WGS-minus-WGA differences in substitution rates for each of the 12 substitution types across the methylation motif units in CS reads. Positive values indicate higher substitution rates in WGS than in WGA reads.

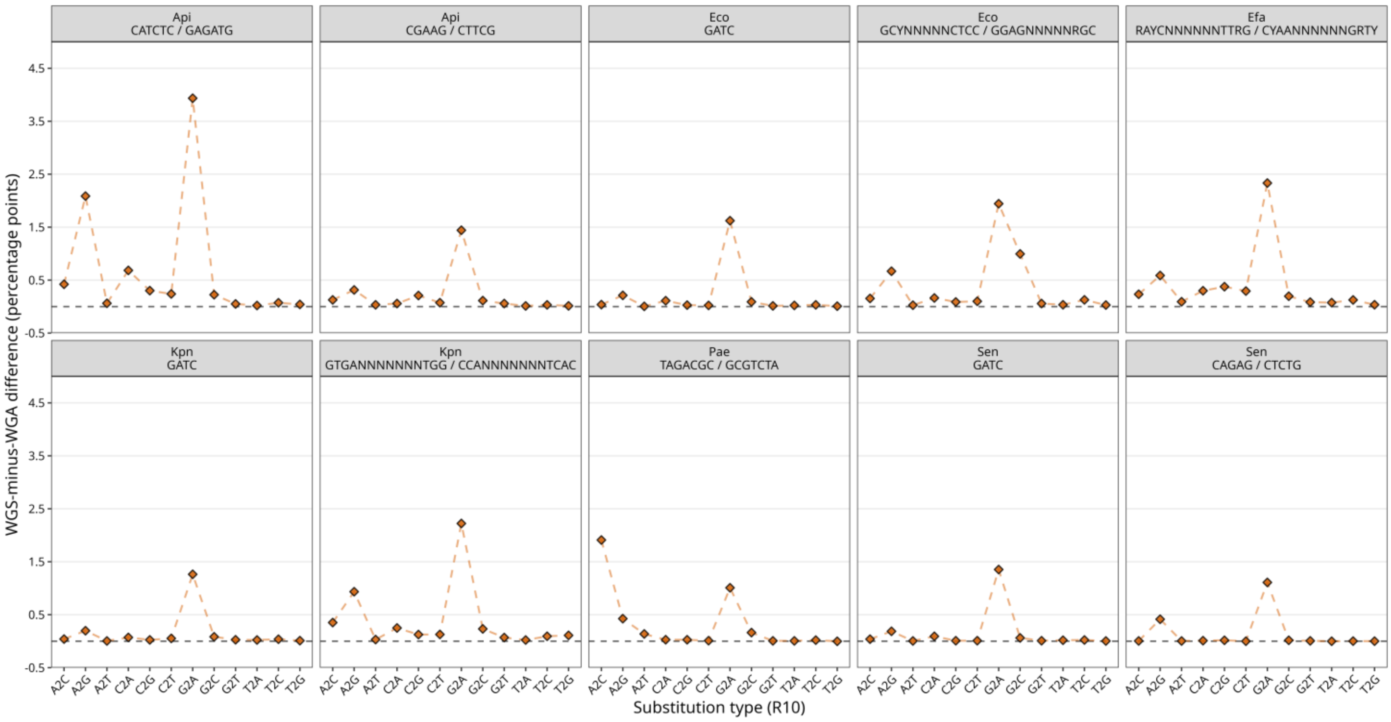


**Figure S11.** WGS-minus-WGA differences in substitution rates for each of the 12 substitution types across the methylation motif units in R10 reads.

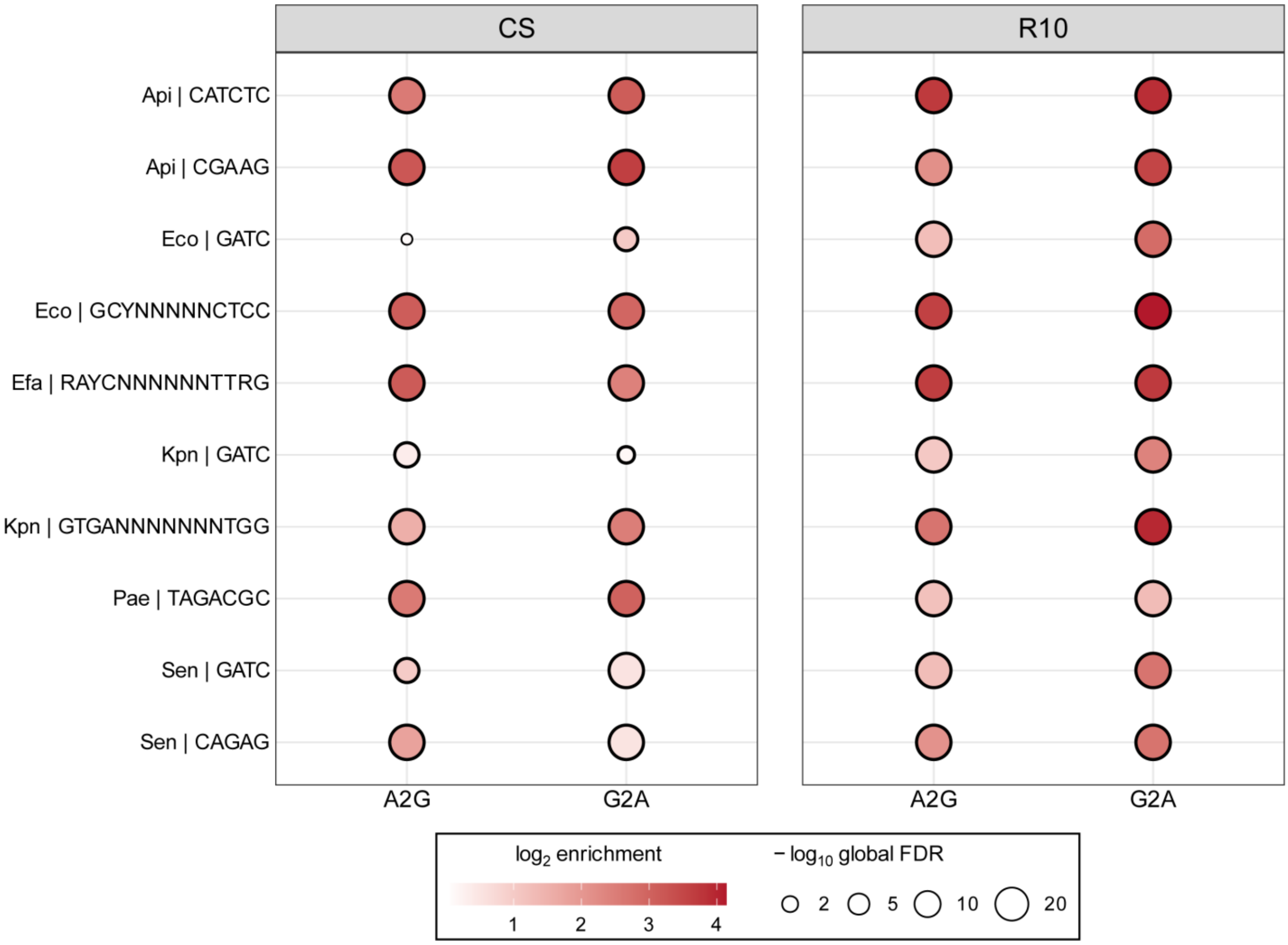


**Figure S12.** Enrichment of motif among genomic sites selected from the top 1% separately for A2G and G2A using the WGS-minus-WGA difference. High-error sites were compared with depth-matched control sites separately for CS and R10. Rows indicate individual species-motif combinations. Color represents the log2 enrichment ratio, with positive values indicating enrichment among high-error sites, and circle size represents -log10 of the globally adjusted false-discovery rate (FDR).

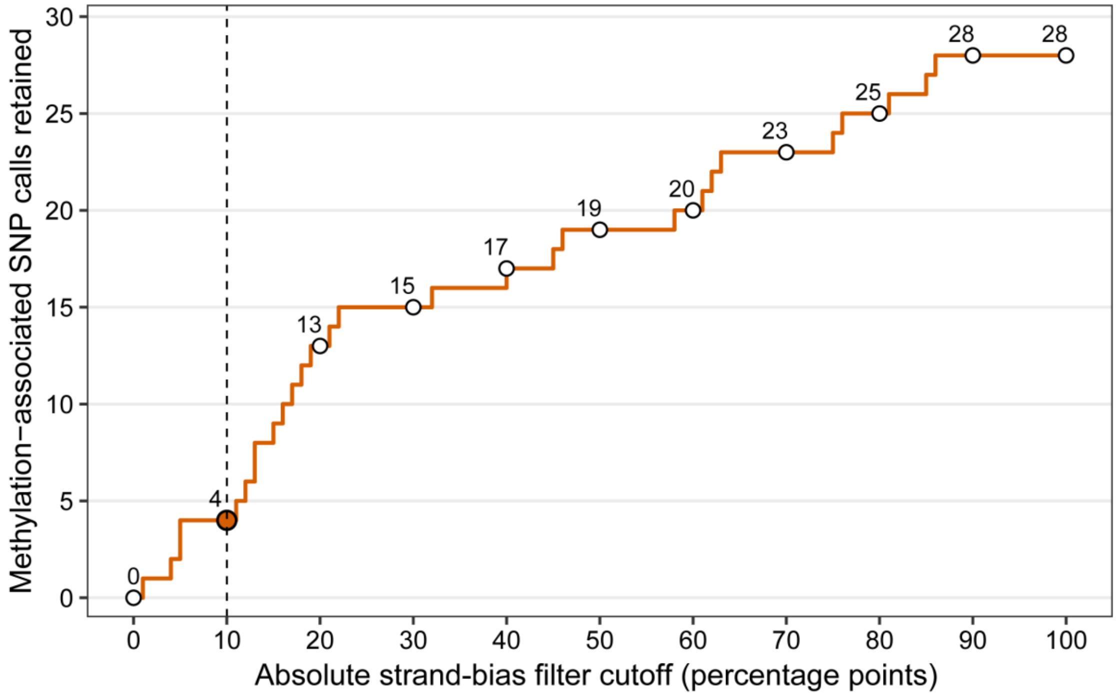


**Figure S13.** Number of the 28 R10 methylation-associated WGS-specific SNP calls remaining after strand-bias filtering across increasing absolute strand-bias thresholds. Strand bias was calculated from DP4 counts as the absolute difference between alternative-allele fractions supported by forward- and reverse-strand reads. Calls meeting or exceeding the specified strand-bias threshold were excluded. The dashed line indicates a cutoff of 10 percentage points, at which four of the 28 candidate calls remained.
